# Prediction of Multi peptides Vaccination from VP10,VP21, VP51 against Reverse Transcriptase Human immunodeficiency Viruses Using Immuno-informatics Approach

**DOI:** 10.1101/2019.12.16.877555

**Authors:** Safa Ahmad Almostafa, Ienas Ibrahim mohmed, Habab Abd elmoneim Siddig, Sahar obi Abd albagi, Nadir Musa Khalil Abuzeid

**Affiliations:** Department of microbiology, MSC of Medical Laboratory Sciences, National University, Sudan; Department of microbiology, faculty of Medical Laboratory Sciences, Alneelain University, Sudan; Associate professor (Omdurman Islamic university) and first Consultant of medical Microbiology)

## Abstract

The human immunodeficiency virus-(HIV) is group of the genus Lentivirus within the family of Retroviridae, subfamily Ortho retrovirinae. Based on genetic characteristics and differences in the viral antigens, HIV is classified into the types 1 and 2 (HIV-1, HIV-2). HIV is identical single – stranded RNA molecule that are enclosed within the core of the virus particle proteins, the genome of the HIV Provirus, also known as DNA, is generated by the Protease against reverse transcriptase RNA genome into DNA, degradation of the RNA and integration of the double – stranded HIV DNA into the human genome. The aim of this study is to determine antigenic peptides from p10, p21, and p51 proteins that can be used for multiple peptide vaccine design using In-Silico study. A total of 73 sequences of three proteins were obtained from NCBI and subjected to multiple sequence alignments using CLUSTALW tool to determine conserved regions.

Immune Epitope Data Base tools were used to determine B cell epitopes, these tools are Bepipred Linear B cell epitopes prediction, surface accessibility and antigenicity prediction. Epitope binding to MHC class I and class II and their population coverage were also determined using IEDB software. The analysis results are as follow, for B cell binding from p10 (^708^**QGYSP**^712^), from p21 (^704^**QGYSP^708^**, ^73^**CVPTDPNPQ**^81^) and (^346^“**FKNL^349^**) from p51. All these peptides have high score in Linear B cell epitopes prediction, surface accessibility and antigenicity prediction. On another hand peptides that reacted to MHC class I were (^47^**EANTTLFCA**^55^, ^53^**FCASDAKAY**^61^, ^55^,**ASDAKAYE**T^63^) form p10,(^38^**YYGVPVWKE**^46^, ^10^**PQEVFLVNV**^18^ and ^29^**AAGSTMGAA**^37^) from p21 and (^63^“**EWEFVNTPP^71^**, **^70^PPLVKLWYQ^78^ and ^79^EKEPIVGA^87^**) from p51 protein respectively. It worth noting that the peptides (^119^**IISLWDQSL***^127^,* ^108^**CVKLTPLCV***^116^*) from p10, (^38^**YYGVPVWKE**^46^, ^20^**LLQYWSQEL**^34^, ^16^**FNMWKNNMV**^30^) from p21 and (**^7^WKGSPAIFQ^21^, ^11^WEFVNTPPL^25^ and ^58^ FLWMGYELH^72^**) protein is also binds to MHC class II with high affinity. All T cell peptides had highest population coverage, and the combined coverage for all peptides in this study was found to be 100%. Using In-Silico studies will ensure less risk of virulence and side effects. Evaluation of antibodies response in animal models is needed to confirm efficacy of these epitopes in inducing protective immune response.

## 1.1. Introduction

The human immunodeficiency virus-(HIV) is group of the genus Lentivirus within the family of Retroviridae, subfamily Ortho retrovirinae (1). Based on genetic characteristics and differences in the viral antigens, HIV is classified into the types 1 and 2 (HIV-1, HIV-2). The immunodeficiency viruses of non-human primates (simian immunodeficiency virus, SIV) are also group of the genus Lentivirus. Epidemiologic and phylogenetic analyses currently available imply that HIV was introduced into the human population around 1920 to 1940. HIV-1 evolved from non-human primate immunodeficiency viruses from Central African chimpanzees (SIVcpz) and HIV-2 from West African sooty mangabeys (SIVsm) (2,3,4).

The HIV genome consists two identical single-stranded RNA molecules that are enclosed within the core of the virus particle. The genome of the HIV provirus, also known as proviral DNA, is generated by the reverse transcription of the viral RNA genome into DNA, degradation of the RNA and integration of the double-stranded HIV DNA into the human genome. The DNA genome is flanked at both ends by LTR (long terminal repeat) sequences. The 5′ LTR region codes for the promotor for transcription of the viral genes. In the direction 5′ to 3′ the reading frame of the *gag* gene follows, encoding the proteins of the outer core membrane (MA, p17), the capsid protein (CA, p24), the nucleocapsid (NC, p7) and a smaller, nucleic acid-stabilizing protein. The *gag* reading frame is followed by the *pol* reading frame coding for the enzymes protease (PR, p12), reverse transcriptase (RT, p51) and RNase H (p15) or RT plus RNase H (together p66) and integrase (IN, p32). Adjacent to the *pol* gene, the *env* reading frame follows from which the two envelope glycoproteins gp120 (surface protein, SU) and gp41 (transmembrane protein, TM) are derived. In addition to the structural proteins, the HIV genome codes for several regulatory proteins: Tat (transactivator protein) and Rev (RNA splicing-regulator) are necessary for the initiation of HIV replication, while the other regulatory proteins Nef (negative regulating factor), Vif (viral infectivity factor), Vpr (virus protein r) and Vpu (virus protein unique) have an impact on viral replication, virus budding and pathogenesis(5, 6).HIV-2 codes for Vpx (virus protein x) instead of Vpu, which is partially responsible for the reduced pathogenicity of HIV-2 (7). The genome structure of the immunodeficiency viruses of chimpanzees (SIVcpand gorillas (SIVgor) is identical to that of HIV- 1(8).

P10 Retroviral proteases are encoded by a part of the pol gene, for example in that of the human immunodeficiency virus (HIV). The protease gene is located between the gag gene (encoding structural proteins) and other enzymatic genes, such as reverse transcriptase and integrase. There are 93 sequences belonging to the retroviral protease family A2 of the aspartic peptidase clan AA at present, according to the Merops database, which provides information on viral as well as other proteases.

Inhibition of HIV-1 replication in monocyte-derived DCs (MDDCs) is associated with an increased expression of p21, a cell cycle regulator that is involved in the differentiation and maturation of DCs. Induction of p21 in MDDCs decreases the pool of dNTPs and increases the antiviral active.

HIV reverse transcriptase (RT) plays a multifunctional role in the transformation of viral RNA into dsDNA and represents a primary target for treatment of AIDS. Currently, all of the drugs in clinical use target the mature RT p66/p51 heterodimer, however, a single p66 peptide chain functions as the precursor for each subunit of the RT heterodimer, requiring a complex maturation process that includes subunit- selective elimination of a single ribonuclease H (RH) domain.

Based on their genetic make-up, HIV-1 viruses are divided into three groups (eg, M [main], N, and O group,). These HIV-1 groups and HIV-2 probably result from distinct cross-species transmission events(9) Pandemic HIV-1 has diversified into at least nine subtypes and many circulating recombinant forms(10, 11) which encode genetic structures from two or more subtypes (eg, A/E=CRF01; A/G=CRF02). The continuously evolving HIV-1 viral diversity poses an immense challenge to the development of any preventive or therapeutic intervention (10). Infection with two or more genetically distinct viruses could lead to new recombinant viruses. Recombination takes place at a higher rate than initially predicted,(11) and circulating recombinant forms account for as much as 20% of infections in some regions (eg, southeast Asia).(12)These findings are in agreement with the occurrence of co-infections with multiple distinct isolates in a close temporal context(13, 14)Further, superinfections in which time points of virus acquisition are months to years apart have been described, although at a much lower frequency than co-infections(13,15,16) Collectively, these observations challenge the assumption that HIV-1 acquisition happens only once with a singular viral strain and that, thereafter, the infected individual is protected from subsequent infections(17) This lack of immunisation has substantial implications for vaccine development. Emerging evidence suggests that clinical progression to AIDS might be more rapid in individuals with dual infections,(18) and encouraging safer sex practices in viraemic HIV-1-infected people might be appropriate to keep recurrent exposure to new viral strains to a minimum.

According to present knowledge, the spread of HIV started at the beginning of the 20th century (4, 19) Zoonotic transmission of SIVcpz from chimpanzees (*Pan troglodytes troglodytes*) occurred for HIV-1 group M and group O around 1920 and for HIV-1 group N around 1960 (20, 21) in West Central Africa. HIV-2 was transmitted zoonotically from sooty mangabey (*Cercocebus atys*) to human in West Africa around 1940 (22) Molecular genetic analyses suggest that HIV-1 was exported to Haiti probably in 1966 and arrived in North America approximately 2 years later (4, 23). Since the mid-1980s the different HIV-1 M subtypes have been spreading, leading to a global pandemic. In contrast to HIV-1, HIV-2 initially remained restricted to West Africa because of its significantly lower infectivity.

After HIV-2 was exported to Portugal and France probably during the mid-1960s, the spreading of HIV-2 with a low prevalence especially in Europe, South America and Asia is documented.

Globally, an estimated 35 million people were living with HIV in 2013 (24). Since 1999 the number of new infections has been decreasing continuously, and for 2013 a number of 1.9 million newly infected persons was estimated. About three quarters of HIV-infected persons live in Sub-Saharan Africa, and also about two-thirds of the reported new infections originate from this region. The countries most affected by HIV with a high prevalence among 15- to 49-year- olds are Swaziland (approximately 27%), Lesotho (approximately 23%) and Botswana (approximately 23%).

An estimated 38·6 (33·4–46·0) million people live with HIV-1 worldwide, while about 25 million have died already.(25)In 2005 alone, there were 4·1 million new HIV-1 infections and 2·8 million AIDS deaths.(25)These estimates mask the dynamic nature of this evolving epidemic in relation to temporal changes, geographic distribution, magnitude, viral diversity, and mode of transmission. Today, there is no region of the world untouched by this pandemic.

Sudan is bordered by countries with high rates of HIV infection. The first AIDS case was reported in 1986, and by 2011 the total number of cases reported had increased to 2218(26). In 2002, a behavioral and epidemiological survey (BES) was carried out by Sudan National AIDS Program [9]. According to the survey, the prevalence among general population was 1.6% (26, 27). Although this survey was the largest survey conducted in Sudan, it was inconclusive largely due to study limitations and mixing of samples that included both general and at-risk populations (27).

The spread of HIV is influenced by poverty and illiteracy, both of which are widespread in Sudan. The movement of people displaced by harsh environmental conditions has contributed to an increase in the number of HIV/AIDS cases. Refugees from other conflicts in the region also flee to Sudan. The war in Darfur resulted in the movement of large population fleeing from the war. This resulted in displacement of about 2.5 million people with many contracted the disease (28). The new war in Blue Nile and South Kordofan also contributed to unfavorable socioeconomic and health situations in Sudan (28).Today, there is no region of the world untouched by this pandemic.

The initial period following the contraction of HIV is called acute HIV, primary HIV or acute retroviral syndrome.(29, 30) Many individuals develop an influenza-like illness or a mononucleosis-like illness 2–4 weeks after exposure while others have no significant symptoms.(31, 32) Symptoms occur in 40–90% of cases and most commonly include fever, large tender lymph nodes, throat inflammation, a rash, headache, tiredness, and/or sores of the mouth and genitals.(29, 32) The rash, which occurs in 20–50% of cases, presents itself on the trunk and is maculopapular, classically.(33) Some people also develop opportunistic infections at this stage.(29) Gastrointestinal symptoms, such as vomiting or diarrhea may occur.(32) Neurological symptoms of peripheral neuropathy or Guillain–Barré syndrome also occurs.(32) The duration of the symptoms varies, but is usually one or twoweeks.(32)

Due to their nonspecific character, these symptoms are not often recognized as signs of HIV infection. Even cases that do get seen by a family doctor or a hospital are often misdiagnosed as one of the many common infectious diseases with overlapping symptoms. Thus, it is recommended that HIV be considered in people presenting with an unexplained fever who may have risk factors for the infection.(32)

The initial symptoms are followed by a stage called clinical latency, asymptomatic HIV, or chronic HIV.(29)Without treatment, this second stage of the natural history of HIV infection can last from about three years(34) to over 20 years(35) on average, about eight years).(36)While typically there are few or no symptoms at first, near the end of this stage many people experience fever, weight loss, gastrointestinal problems and muscle pains. Between 50 and 70% of people also develop persistent generalized lymphadenopathy, characterized by unexplained, non-painful enlargement of more than one group of lymph nodes (other than in the groin) for over three to six months(37).

## Materials and Methods

### In-Silico, Study Materials and Methods

A total of 10 HIV structural poly protein sequences were obtained from NCBI(www.ncbi.nlm.nih.gov/protein/?term=) in FASTA format in Jun 2019 (Table 1) For VP10.

**Table 1:**
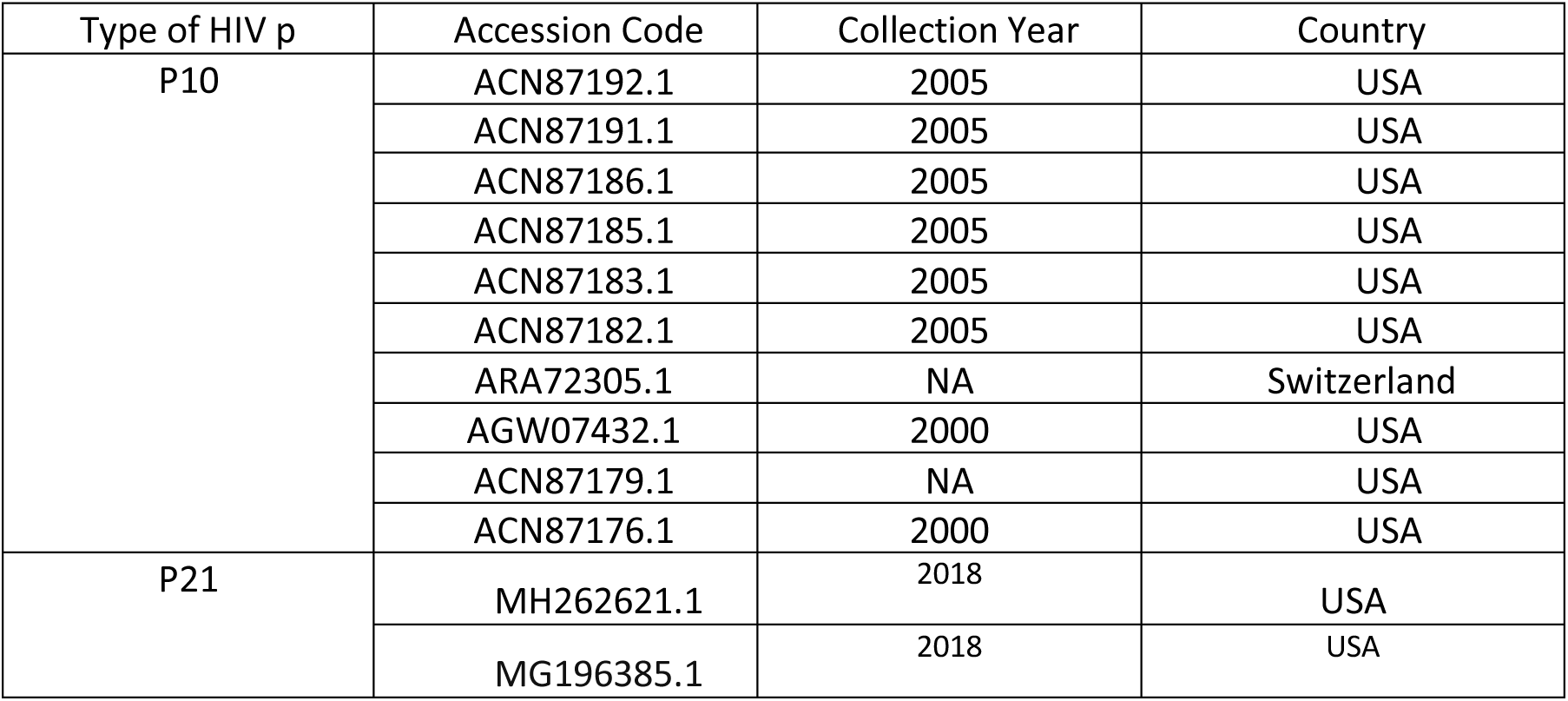
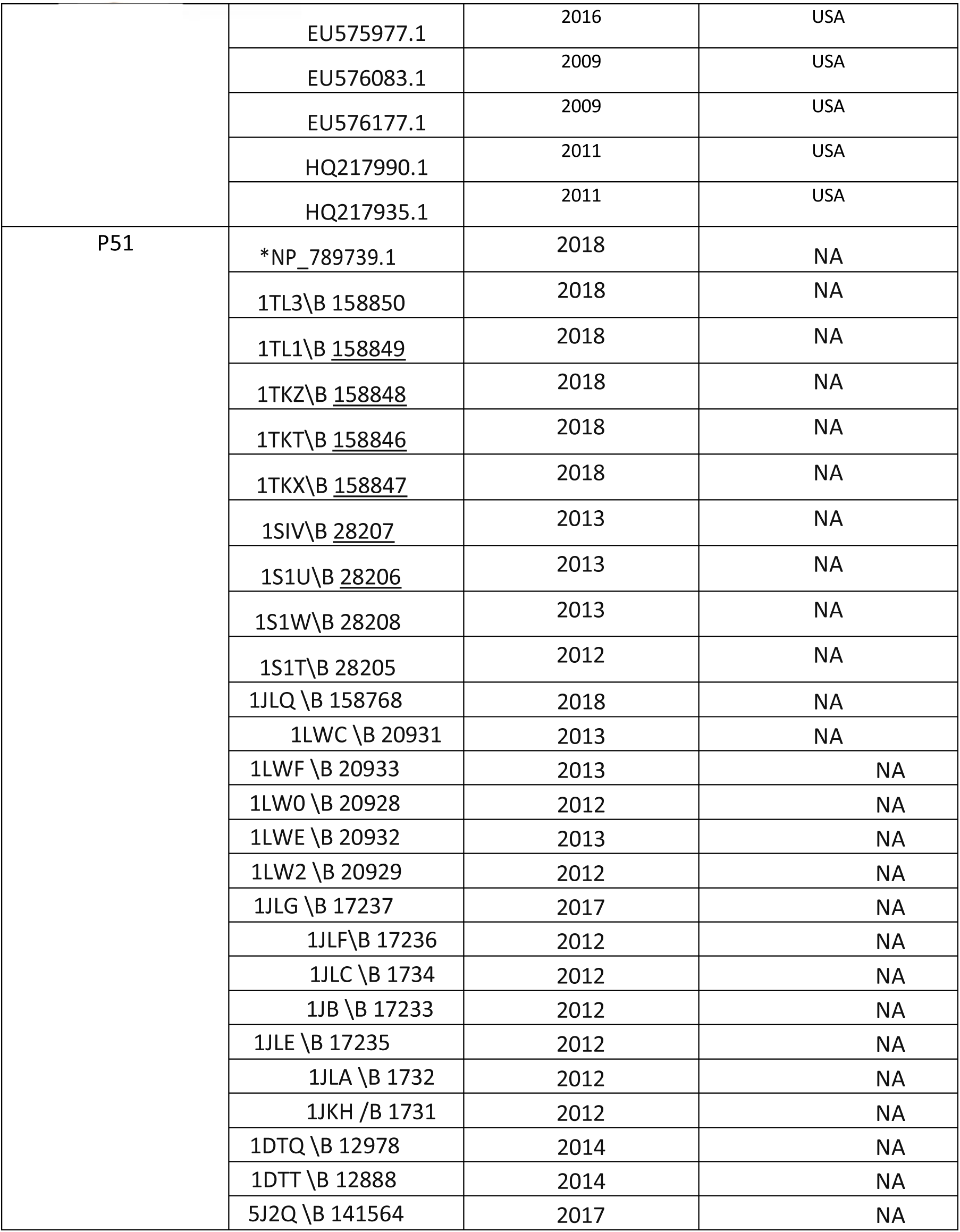
HIV Countries and accession numbers of retrieved sequence

Total of 6 HIV structural poly protein sequences were obtained from NCBI (www.ncbi.nlm.nih.gov/protein/?term=) in FASTA format in May 2019 For vp21., reference sequences were identified from NCBI reference sequence.

Total of 57 reverse transcriptase protein (RT P51) sequences of HIV virus were retrieved from the National Center for Biotechnology Information database (NCBI) (www.ncbi.nlm.nih.gov/protein/?term=) in May 2019, reference sequences were identified from NCBI reference sequence.

All sequences of structural poly protein were subjected to multiple sequence alignments using CLUSTALW tool of BIOEDIT sequence alignment editor (version 7.2.5.0) in order to identify conserved regions between sequences. Then epitopes prediction and analysis of each protein was done using different tools of immune epitope data base IEDB software (http://www.iedb.org).

#### Determination of conserve sequence and phylogenic analysis

The retrieved sequences were aligned by multiple sequence alignment (MSA) using Bio Edit software (version 7.2.5.0) (38), to obtain the conserved regions. Which were considered candidate epitopes were analyzed by different prediction tools at the Immune Epitope Database IEDB analysis resource (http://www.iedb.org/).

#### Prediction of B cell epitopes

Candidate epitopes were analyzed by several B-cell prediction methods which determine the antigenicity, hydrophilicity, flexibility and surface accessibility (39). The method for predicting B-cell epitopes is the hidden Markov model. The linear predicted epitopes were obtained by using BepiPred linear epitope prediction with a threshold value of -0.165 (40). The surface accessible epitopes were predicted from the conserved region with a threshold value of 1.000 using the Emini surface accessibility prediction tool (41). Using the kolaskar antigenicity we determined the antigenic sites with a threshold value of 1.000 (42).

#### MHC epitope prediction

##### MHC class I epitope prediction

MHC I prediction tool has been used to analyze the referential (p10,p21,P51) RT protein of the HIV binding to MHC class I molecules (http://tools.iedb.org/mhci/). This prediction tool consists of 9 different prediction methods available in the Immune Epitope Database (IEDB) namely the covariance matrix SMMPMBEC, Artificial Neural Network (ANN), Stablized Materix Method (SMM), CombLib-Sidney2008, Consensus, NetMHCpan, PickPocket, NetMHCcons and IEDB Recommended.

Artificial neural network (ANN) was chosen as it is perfectly suitable in the incorporated such higher order connections when predicting the binding affinity (43, 44). Analysis was done using HLA alleles (45). For all the alleles, peptide length was set to 9 amino acids prior to the prediction. The half maximal inhibitory concentration (IC50) values of the peptide’s binding to MHC I molecules was calculated and the ones that had binding affinity (IC50) less than 100 nM were suggested to be promising candidate epitopes.

### MHC Class II binding predictions

The referential protein sequence was submitted in the (IEDB) MHC II binding prediction tool (http://tools.immuneepitope.org/mhcii/). For the screening of promising epitopes, human allele reference set (HLA DR, DQ, DP) was used (46, 47). In MHC II binding analysis at the (IEDB) there are seven prediction methods including IEDB recommended, Consensus, Net MHCII pan, NN-align, SMM-align, combinatorial library and Sturniolo methods. NN-align method was used with a half maximal inhibitory concentration (IC50) 0f 1000 nM. Peptides less or equal to the (IC50) value were proposed to be promising MHC II epitopes (48, 49).

#### Population coverage

The estimation of the population coverage are based on the MHC binding with or without T cell restriction data, there for Nemours based tool was developed to predict population coverage of T cell epitope-based diagnostic and vaccines based on MHC binding with or without T cell restriction data (50). All alleles that interact with epitopes from P51 RT protein were subjected population coverage tool of IEDB (http://tools.iedb.org/tools/population/iedb_input) to calculate the whole world population coverage of MHC class I, MHC II and combined MHC I and II alleles for each protein (51).

#### Homology modeling

**Raptor X** protein structure prediction server available at (http://raptorx.uchicago.edu/StructurePrediction/predict/) was utilized for the creation of the **3D structure** and **Chimera 1.8** (52) was the tool used to visualize the selected epitopes belonging to the B cell, MHC class I and MHC class II.

## Results

### Alignment and determination of conserved regions

A total of10 HIV p10,6 of p21,and 57 HIV p51 sequences were obtained from NCBI (www.ncbi.nlm.nih.gov/protein/?term=) in FASTA format in May 2019., reference sequences were identified from NCBI reference sequence. All sequences of HIVp10,p21 and P51 were subjected to multiple sequence alignments using CLUSTALW tool of BIOEDIT sequence alignment editor in order to identify conserved regions between sequences. Then epitopes prediction and analysis of each protein was done using different tools of immune epitope data base IEDP software (http://www.iedb.org).

**Figure 1:**
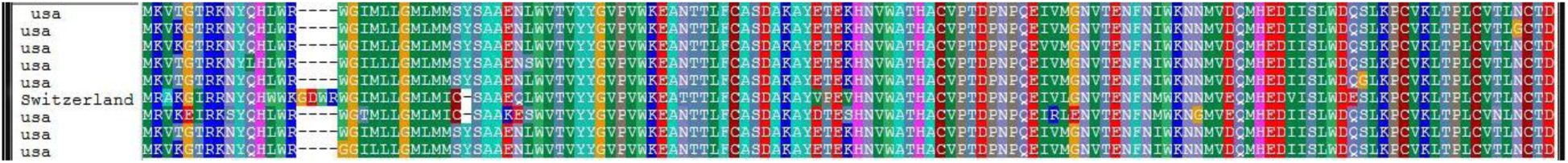
Multiple sequence alignment p10

**Figure 2:**
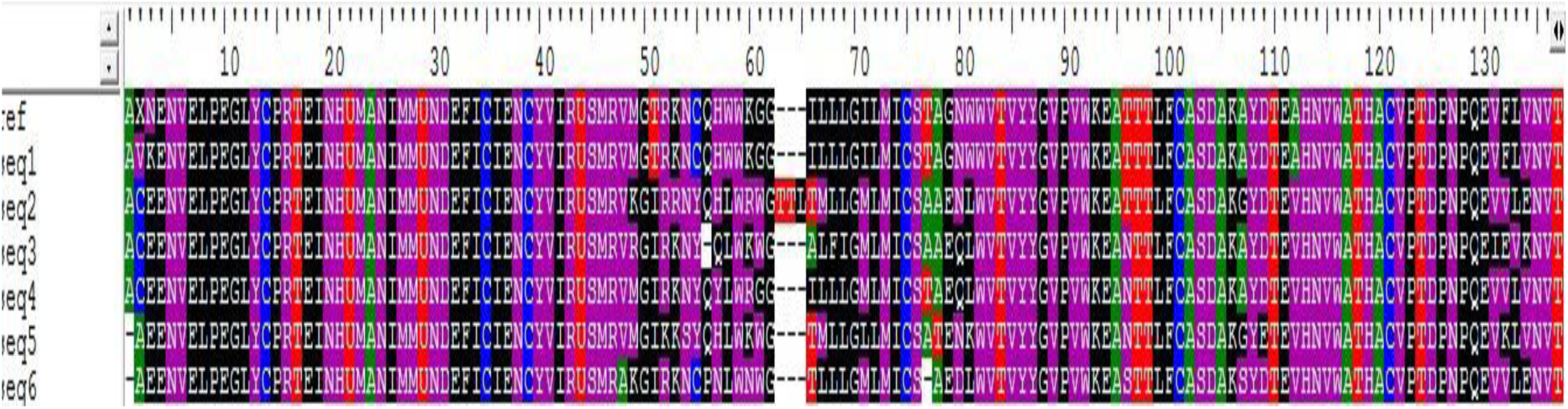
Multiple sequence alignment;p21

**Figure 3:**
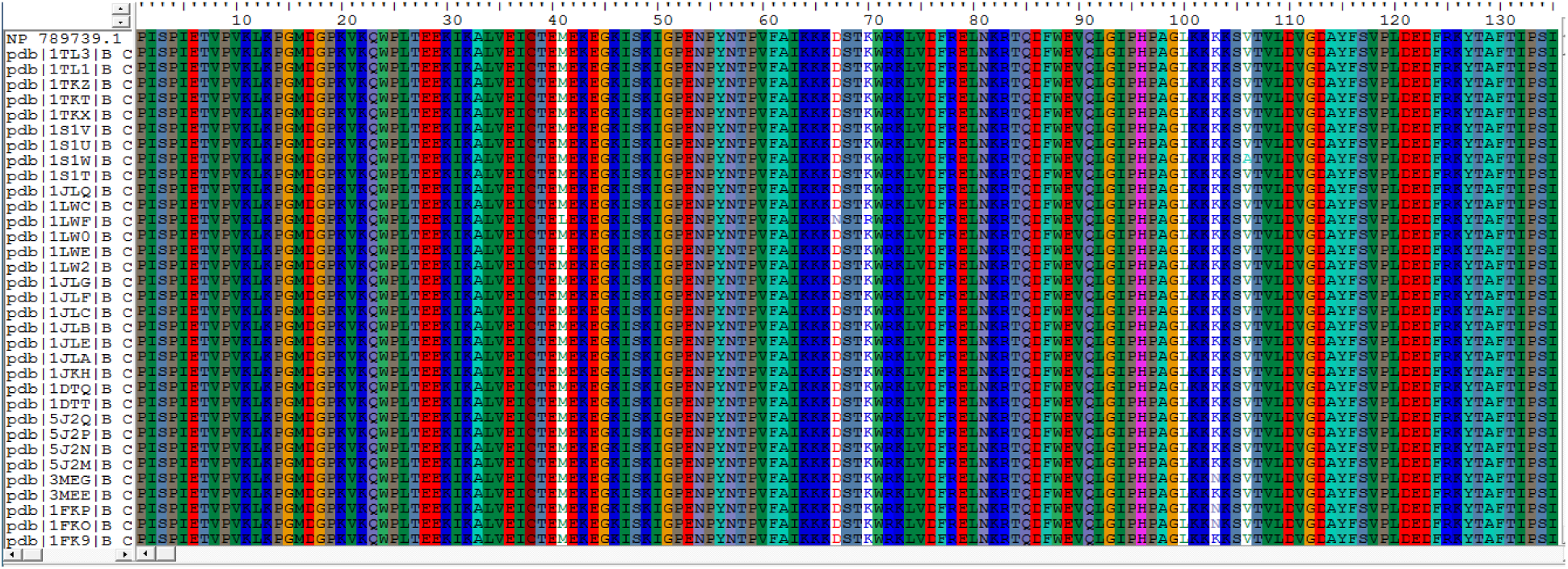
Multiple sequence alignment;p51

#### Prediction of B cell epitopes

Poly structural protein was tested using Bepipred linear epitope production, Emini surface accessibility and Kolaskar and Tongaonkar antigenicity tool of IEDB. All values equal or greater than the default threshold were considered as potential B-cell binders.

Regarding Emini surface accessibility prediction, the average binding score of poly structural protein was 1.00, and minimum and maximum values as displayed in Table 2. All values equal or greater than the default threshold were predicted to have good surface accessibility. All proteins were subjected to Kolaskar and Tongaonkar antigenicity prediction tool of IEDB to predict peptides with probability of being antigenic, the average threshold value for poly structural protein was 1.01. Epitopes with values equal or greater than the average score are considered as antigenic peptides.

**Table 2:**
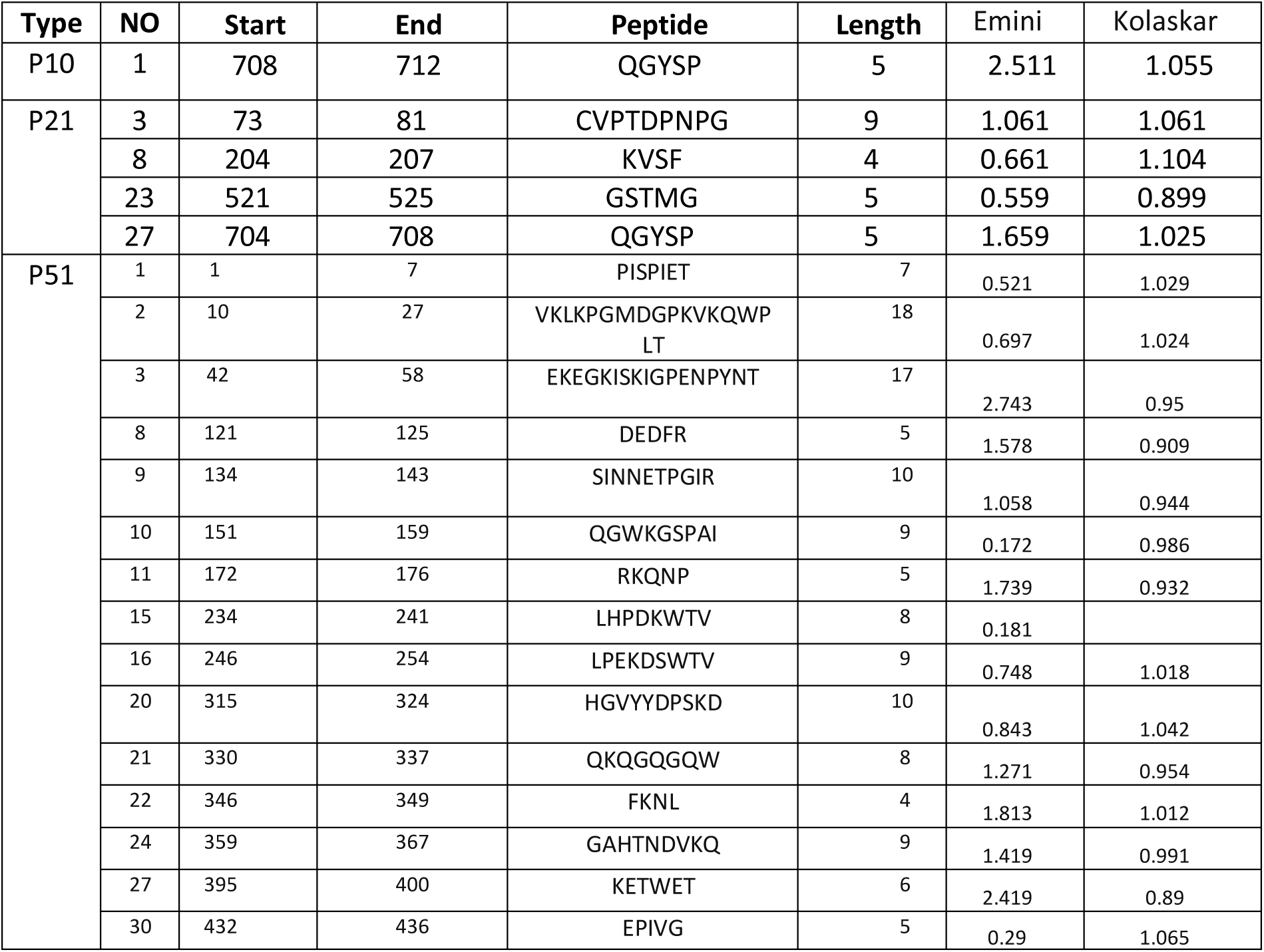
predicted B cell epitopes

The result of structural polyprotein showed that three peptides had passed the antigenicity prediction and surface accessibility prediction test, QGYSP from 704 to 708 was found to have greatest score among all other peptides in both tests in p10 and p21,and one peptides had passed the antigenicity prediction and surface accessibility prediction test, **FKNL** from 346 to 349 was found to have greatest score among all other peptides in both tests in p51. The result is summarized in Table 2.

**Figure 2:**
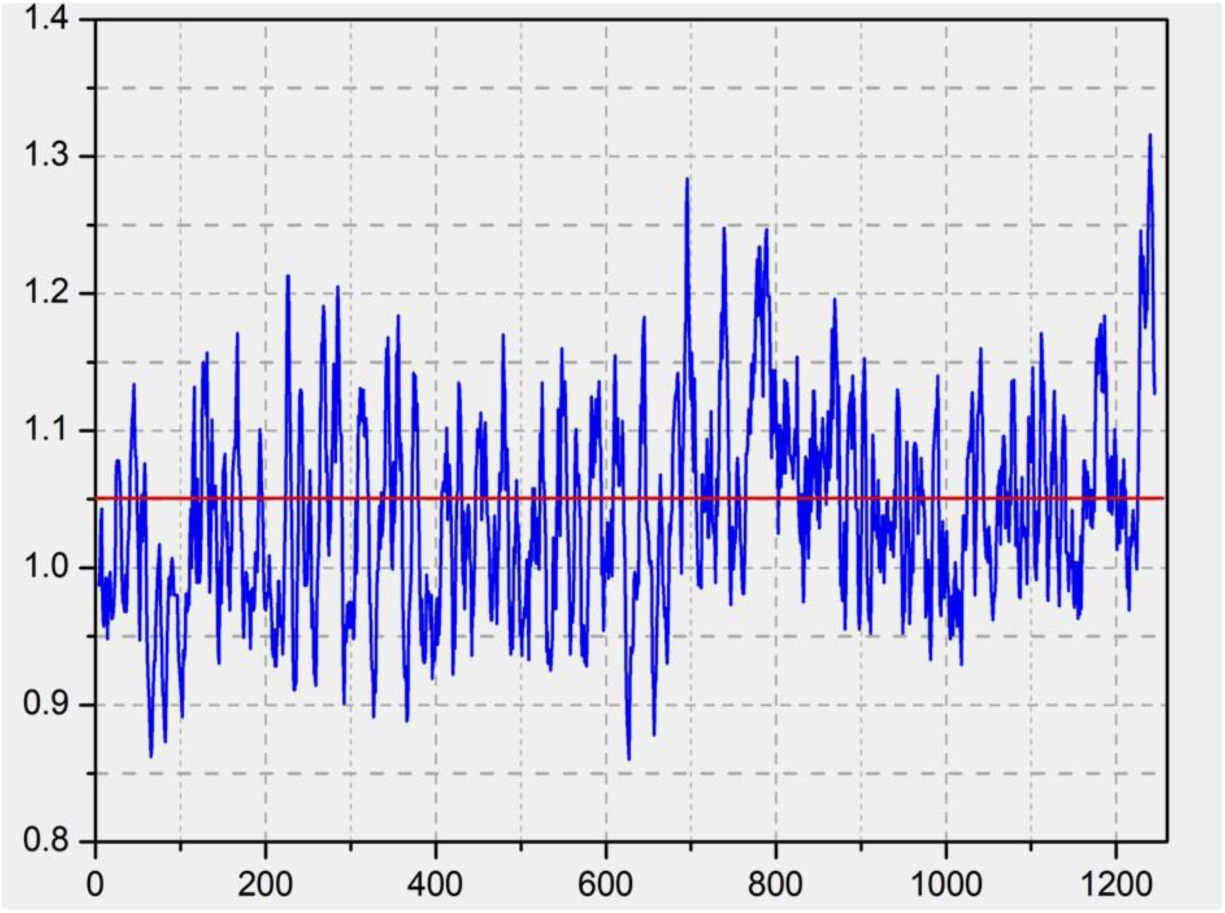
surface accessibility analyses using Emini surface accessibility scale

**Figure 3.3.**
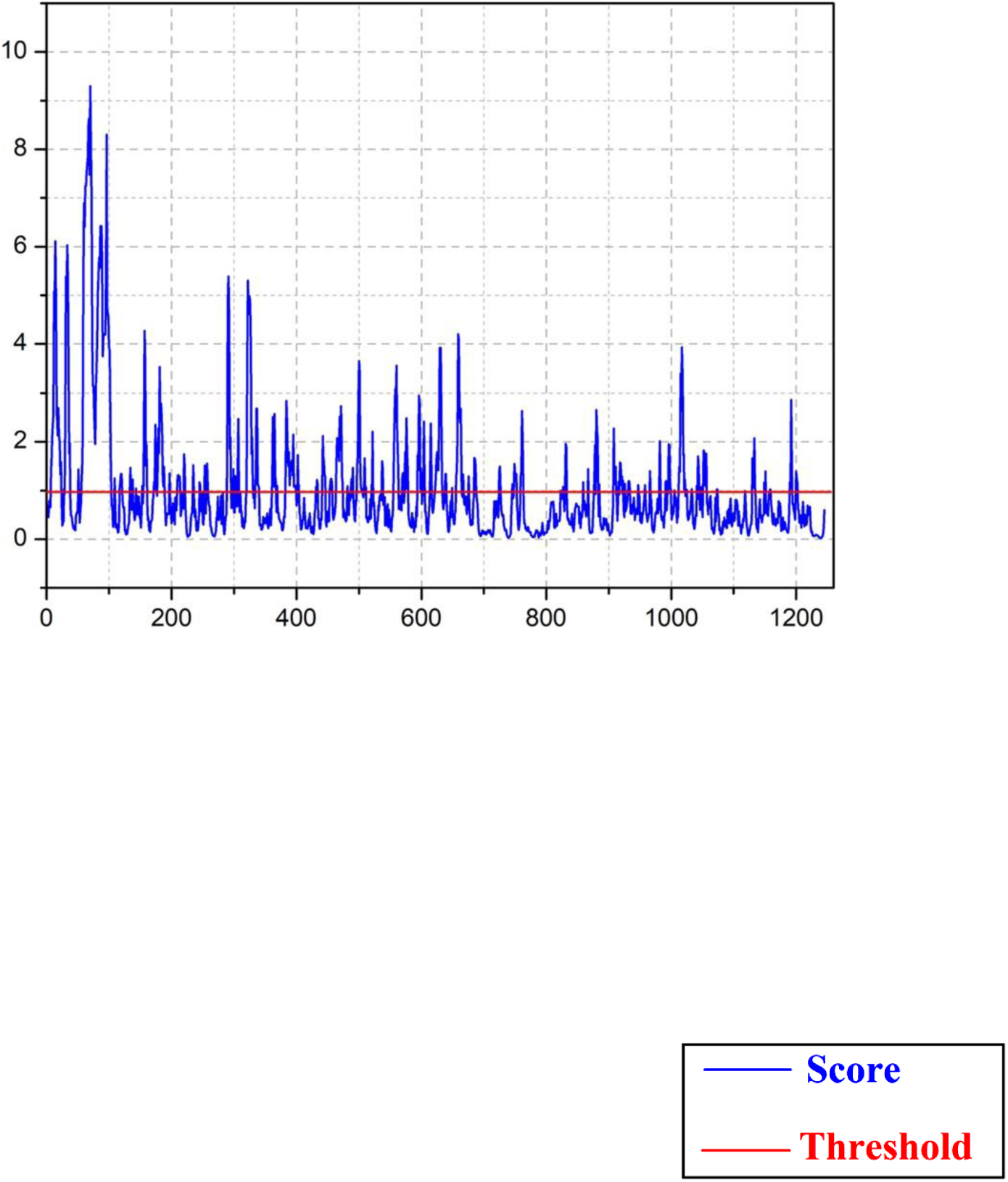
Kolaskar and Tongaonker antigenicit

**Figure 4:**
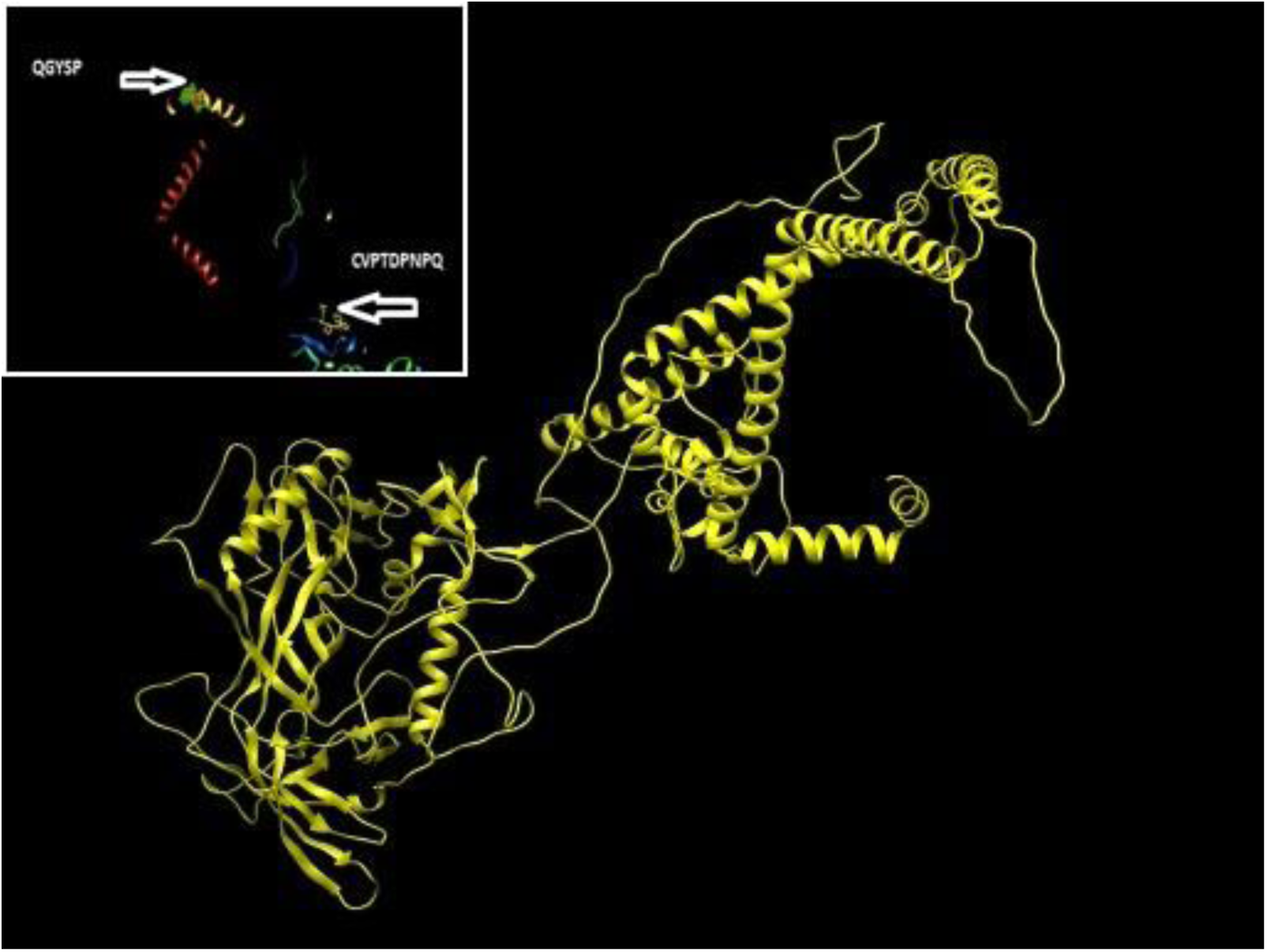
prediction of Bcell epitopes p10 by chimera 1.8

**Figure 5:**
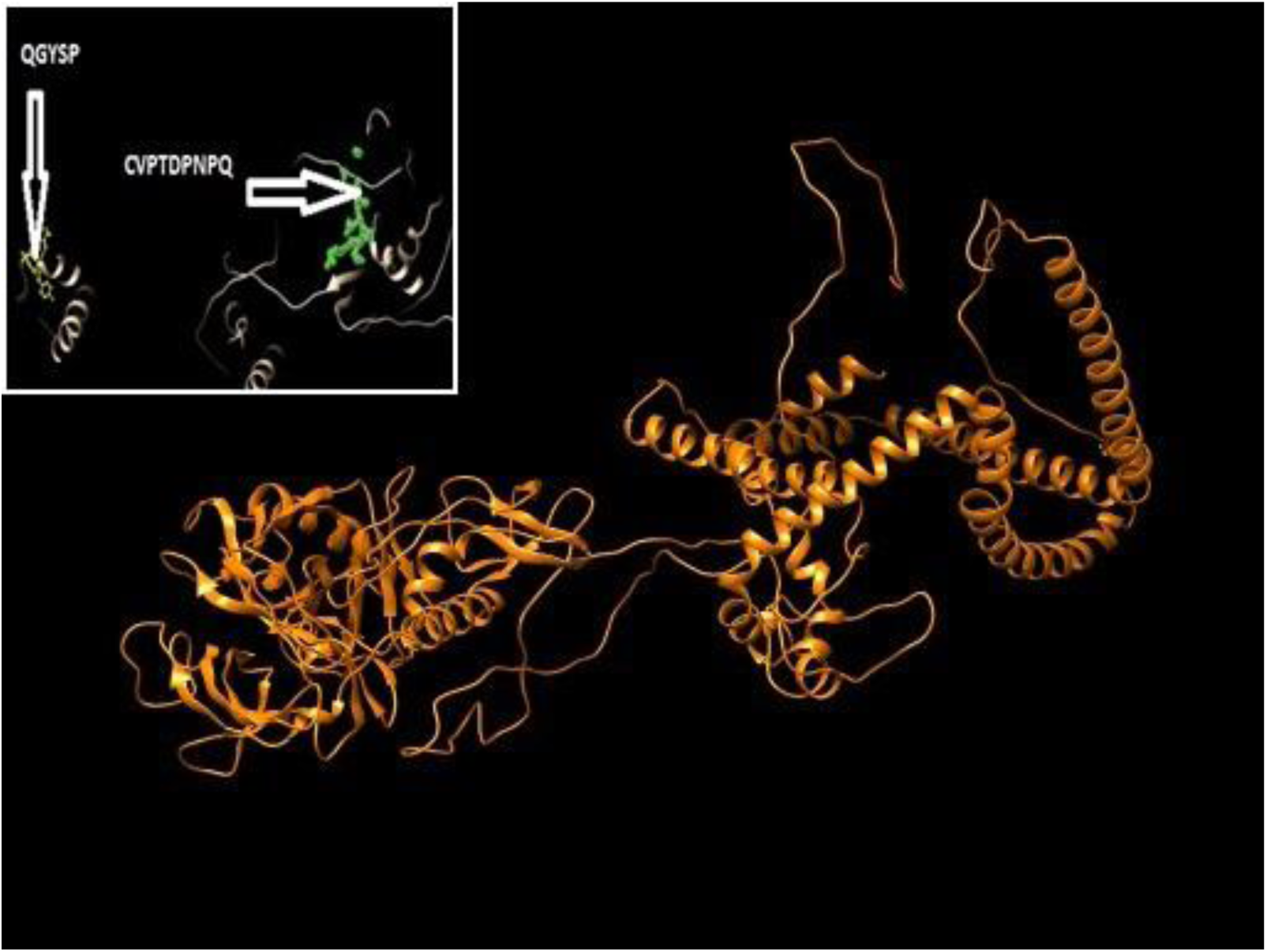
prediction of Bcell epitopes p21 by chimera 1.8

**Figure 6:**
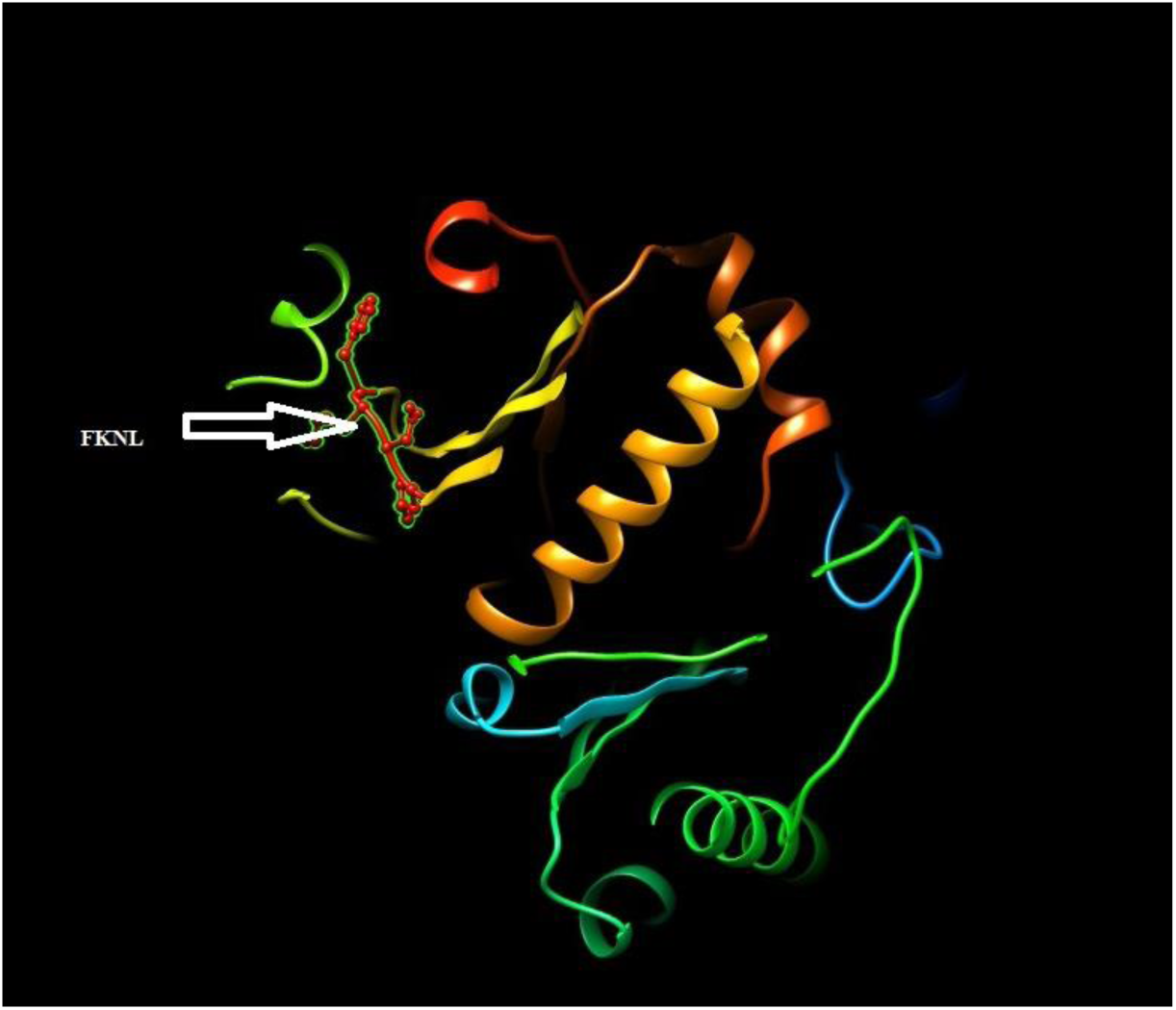
prediction of Bcell epitopes p51 by chimera 1.8

#### Predictions of T cell Epitopes

IEDP server (http://www.iedb.org)was used through specific tools to determine MHC1 and MHC II binding epitopes. This server uses specific scoring IC50 (inhibitory concentration 50) to predict epitopes that bind to different MHC class I and MHC class II alleles.

### Prediction of T-cell epitopes and MHC class I interaction analysis

Epitopes sequences of structural poly protein subjected to MHC class I binding prediction tool of IEDP. T-cell epitope predicted to interact with different MHC class I alleles using ANN (artificial natural network) as prediction method and length of nine amino acids.

Using same score for all protein of 100 IC50, The peptide ^47^EANTTLFCA^55^, ^53^FCASDAKAY^61^, ^55^ASDAKAYET^63^height affinity to interact with the largest number of (6) alleles for p10,12 peptide of **structural polyprotein** were found to interact with MHC class I. The peptide (^38^ YYGVPVWKE^46^,^10^ PQEVFLVNV^18^ and ^29^AAGSTMGAA^37^) had height affinity to interact with the largest number of alleles for p21.

These 10 sequenced epitopes were obtained from the IEDB MHC I binding epitope prediction tool using ANN method and the reference reverse transcripts protein (p51) as an input. There were 44 human MHC class-I alleles for which the epitopes were predicted the best 3 are^63^“**EWEFVNTPP^71^, ^70^PPLVKLWYQ^78^ and ^79^EKEPIVGA ^87^**.

**Table 3:**
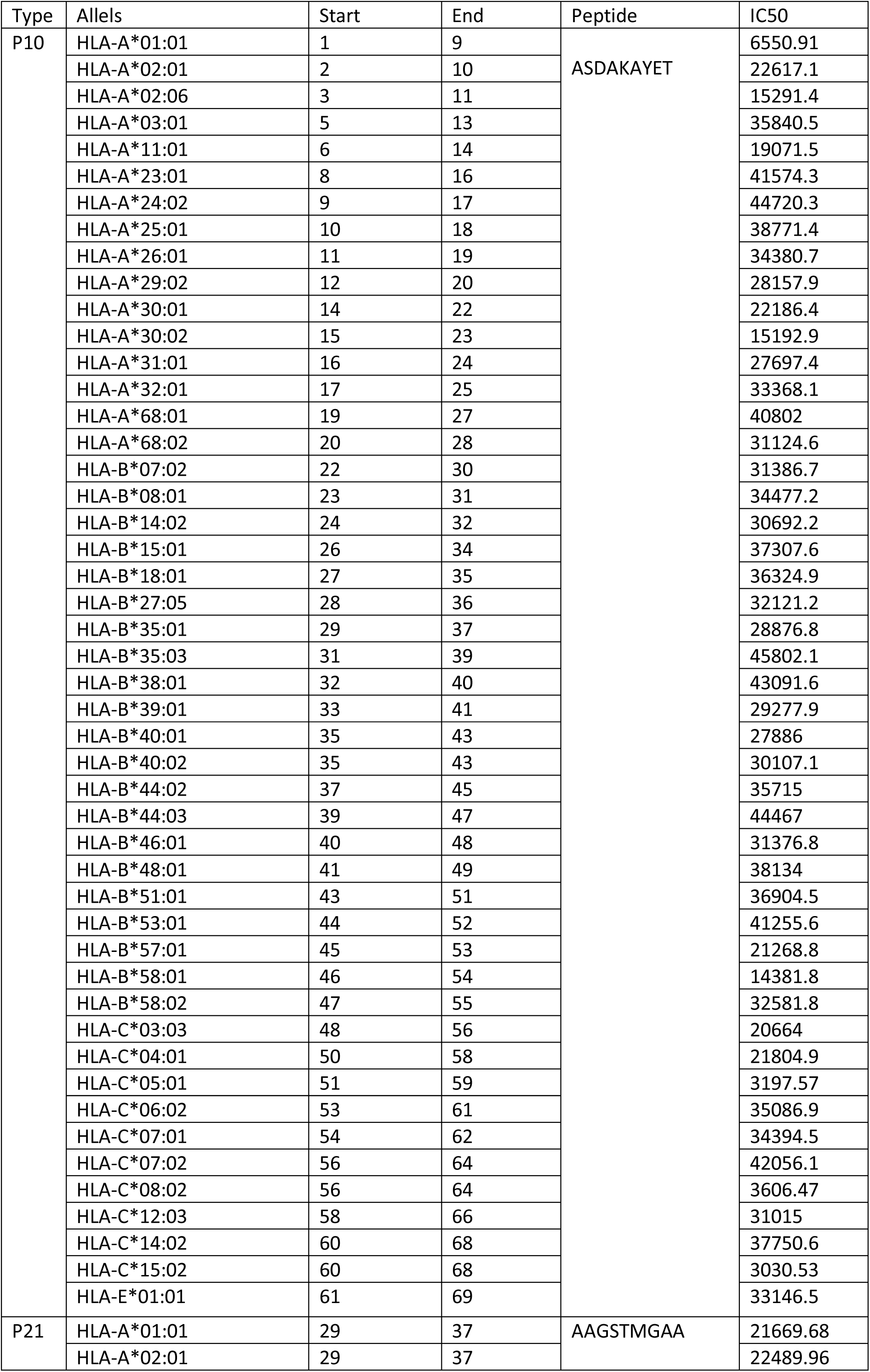
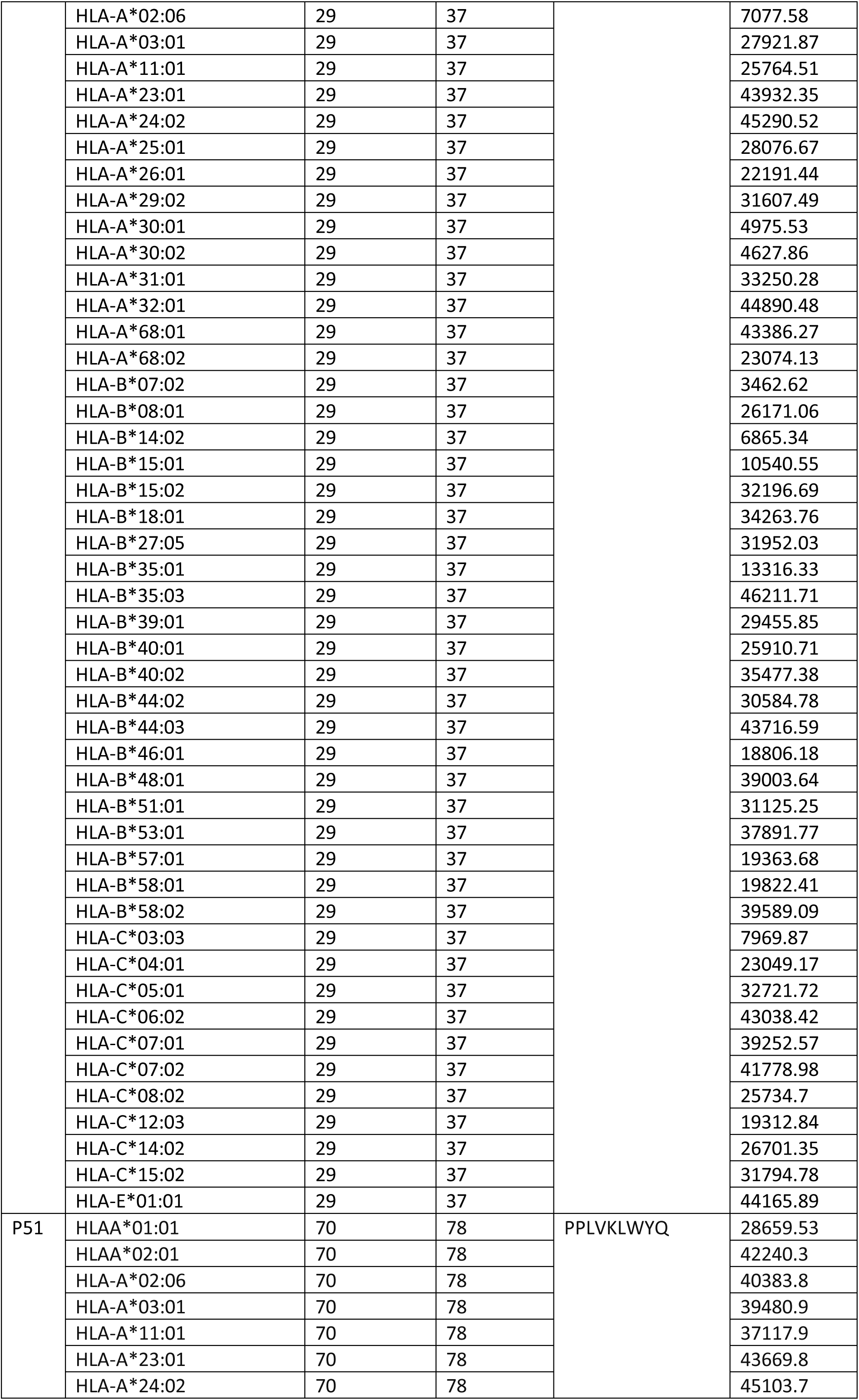
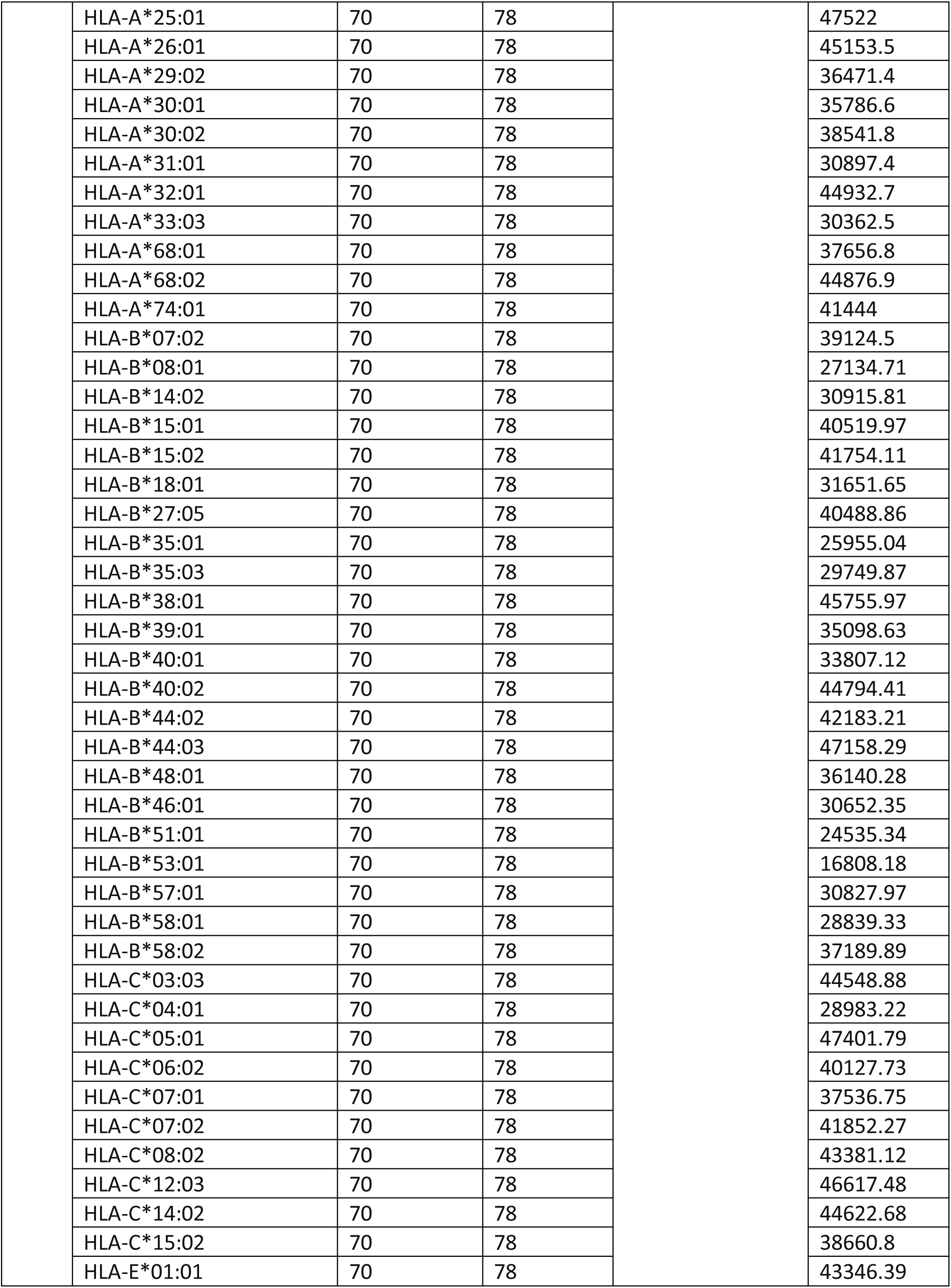
T cell MHC I

**Figure 7:**
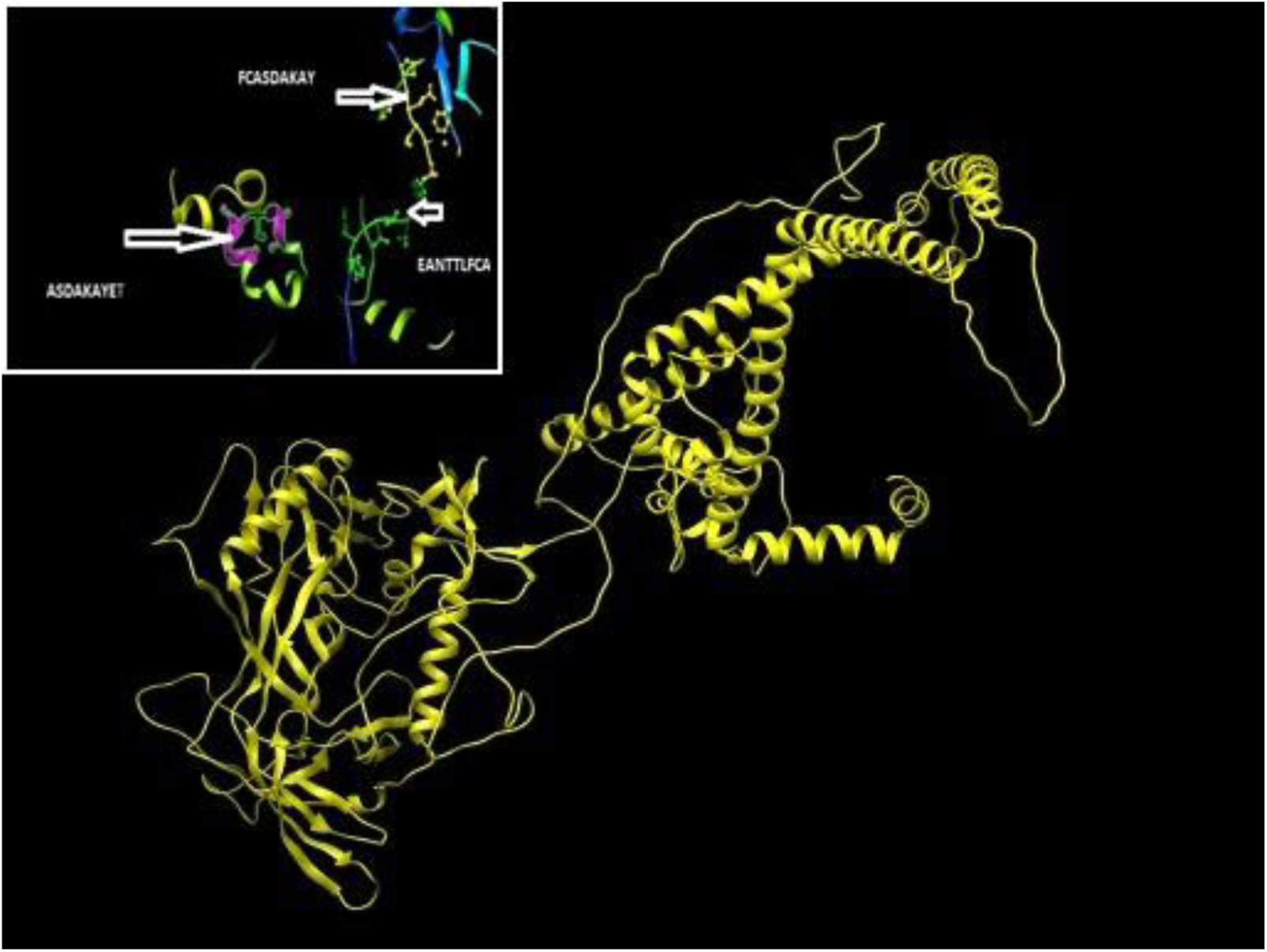
prediction peptides with MHC I of p10 by chimera 1.8

**Figure 8:**
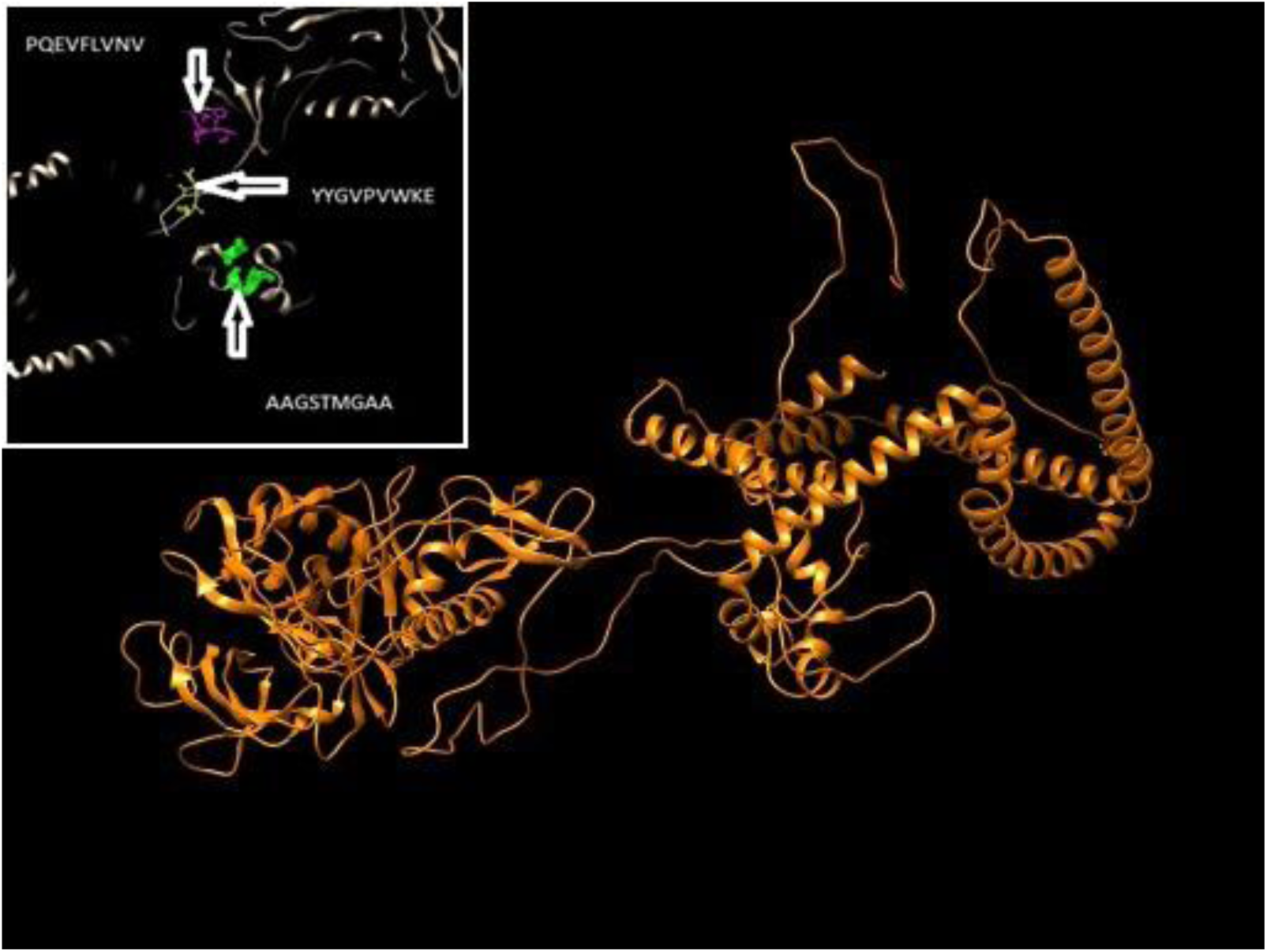
prediction peptides with MHC I of p21 by chimera 1.8

**Figure 9:**
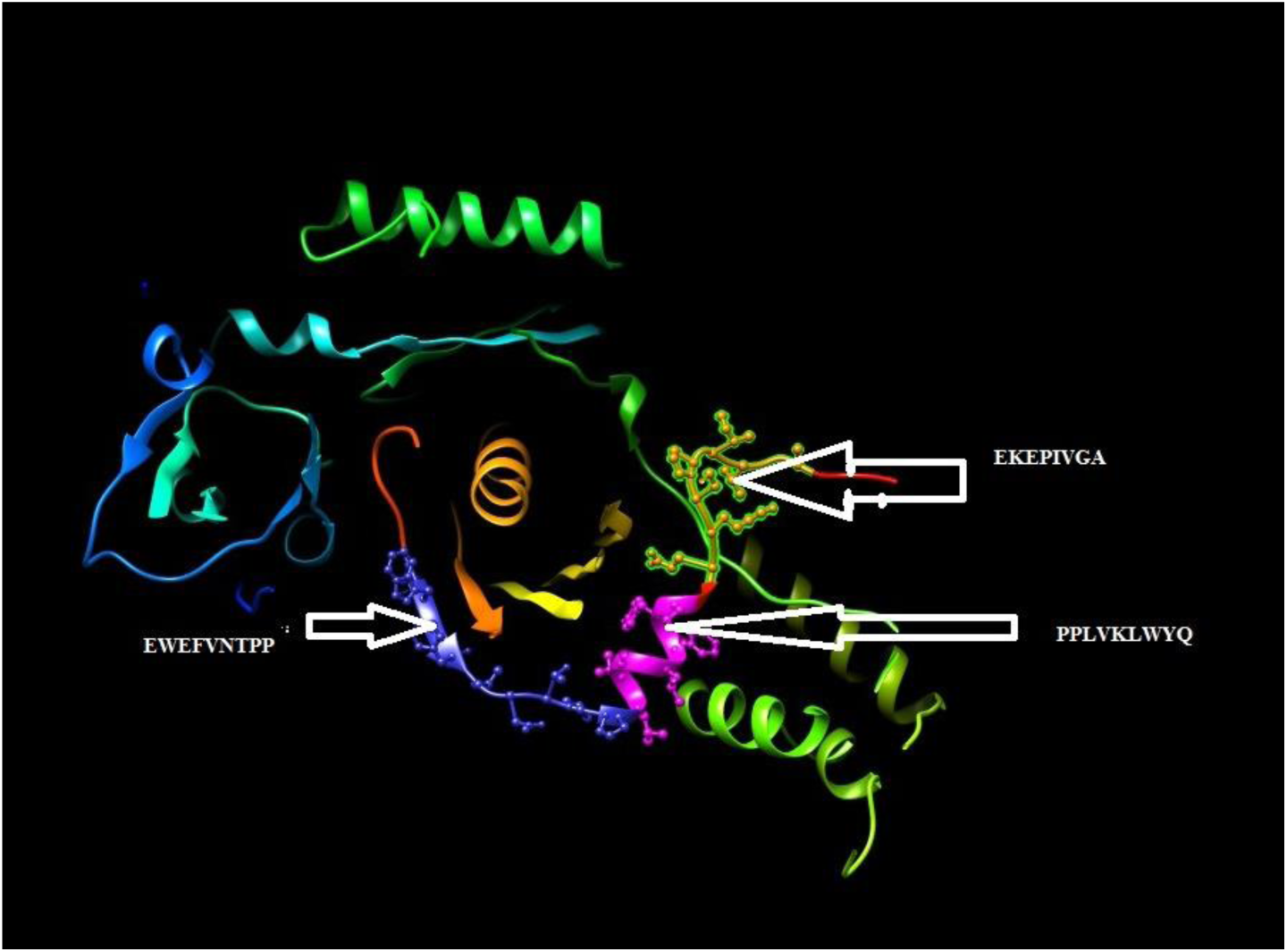
prediction peptides with MHC I of p51 by chimera 1.8

### Prediction of T Helper cell epitopes and interaction with MHC class II

Using MHC class II binding prediction method of IEDB and based on NN align and IC50 of 500 for structural polyprotein. From all predicted epitopes of **structural polyprotein** that bind to MHC class II, the core sequence^119^**IISLWDQSL***^127^,*^108^**CVKLTPLCV***^116^*bind to(25) alleles. which is largest number of alleles in contrast to other predicted, core sequences in p10. the core sequence ^38^**YYGVPVWKE**^52^, ^17^**FNMWKNNMV**^31^, ^20^**LLQYWSQEL**^34^ bind to 43 alleles. which is largest number of alleles in contrast to other predicted, core sequences in p21. the core sequence **^7^WKGSPAIFQ^21^** that showed affinity for 27 MHC- II molecules, ^11^**FLWMGYEL**^25^ that showed affinity for 43 MHC- II molecules, and ^58^**WEFVNTPPL^72^** that showed affinity for 37 MHC- II molecules.

**Table 4:**
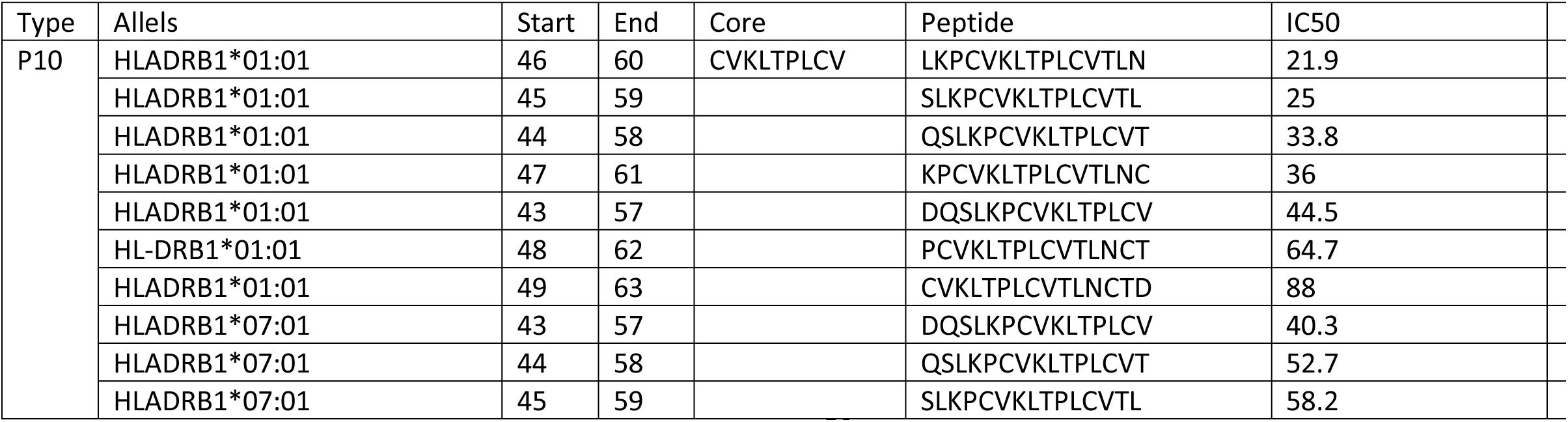
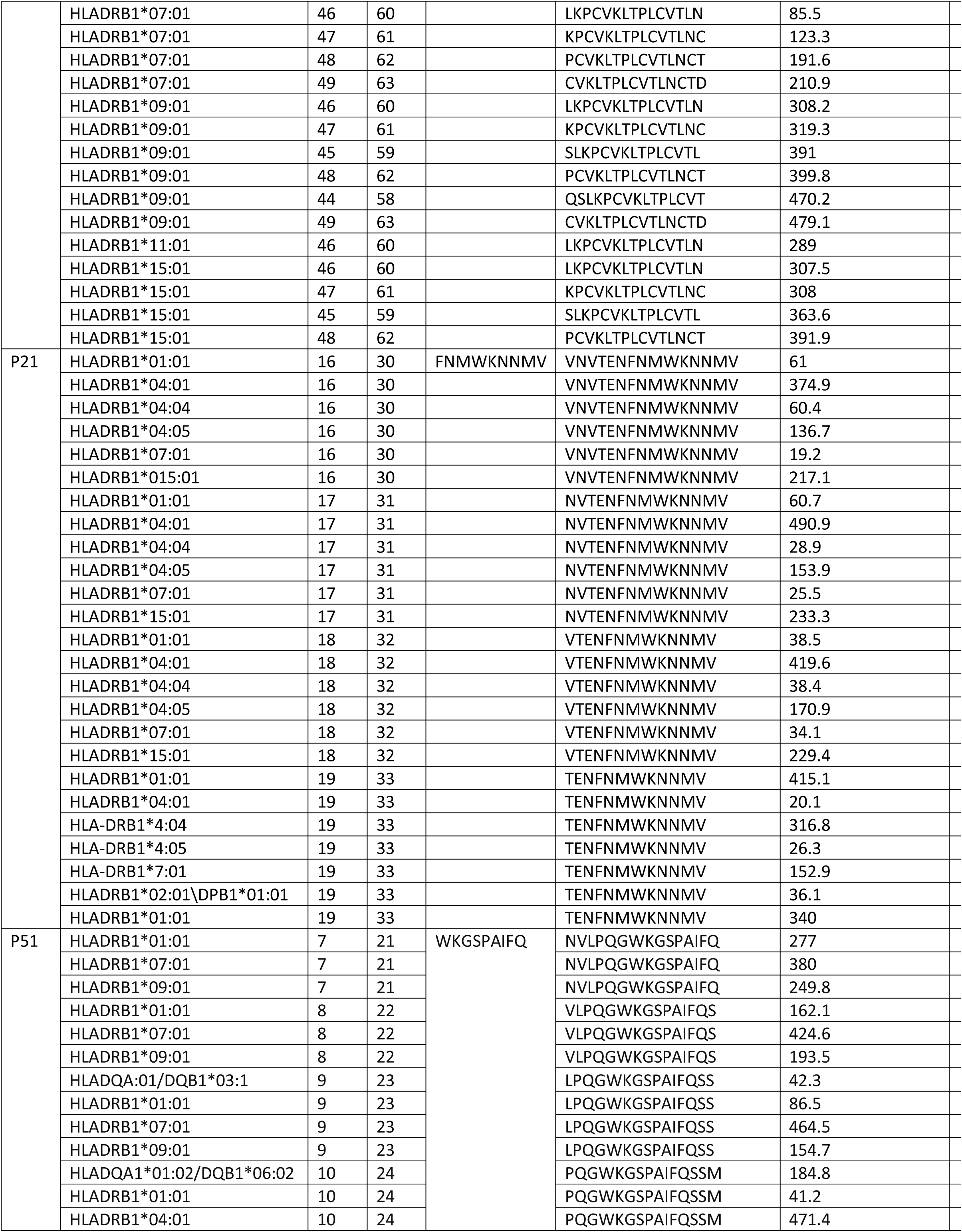
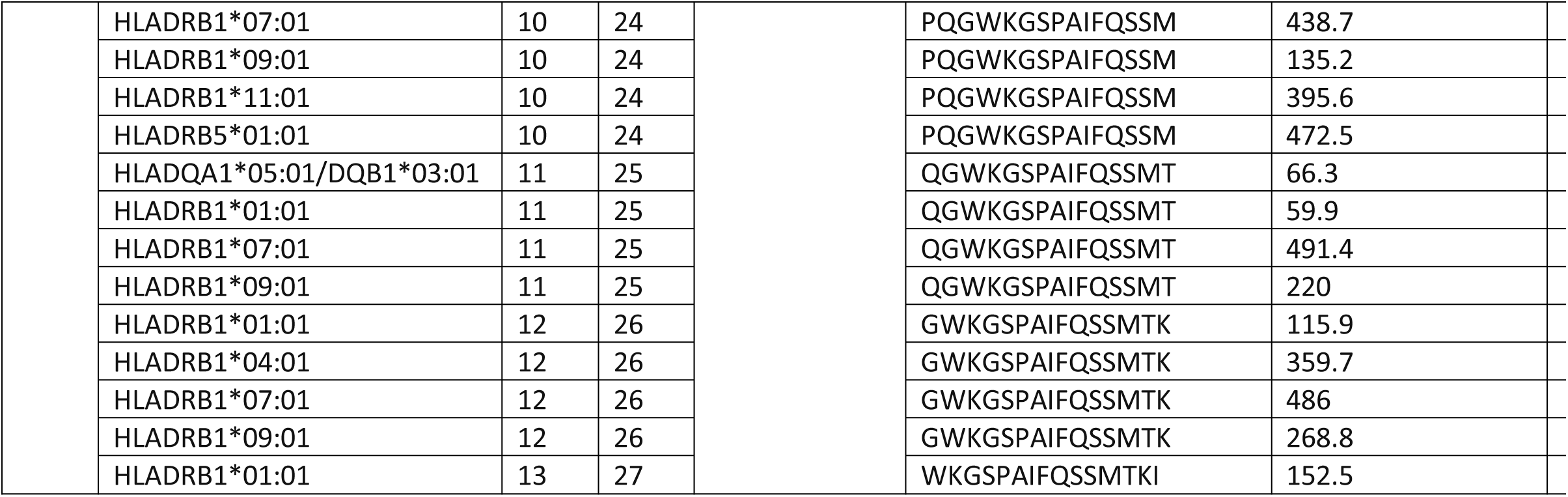
T cell MHC II of HIV

**Figure 10:**
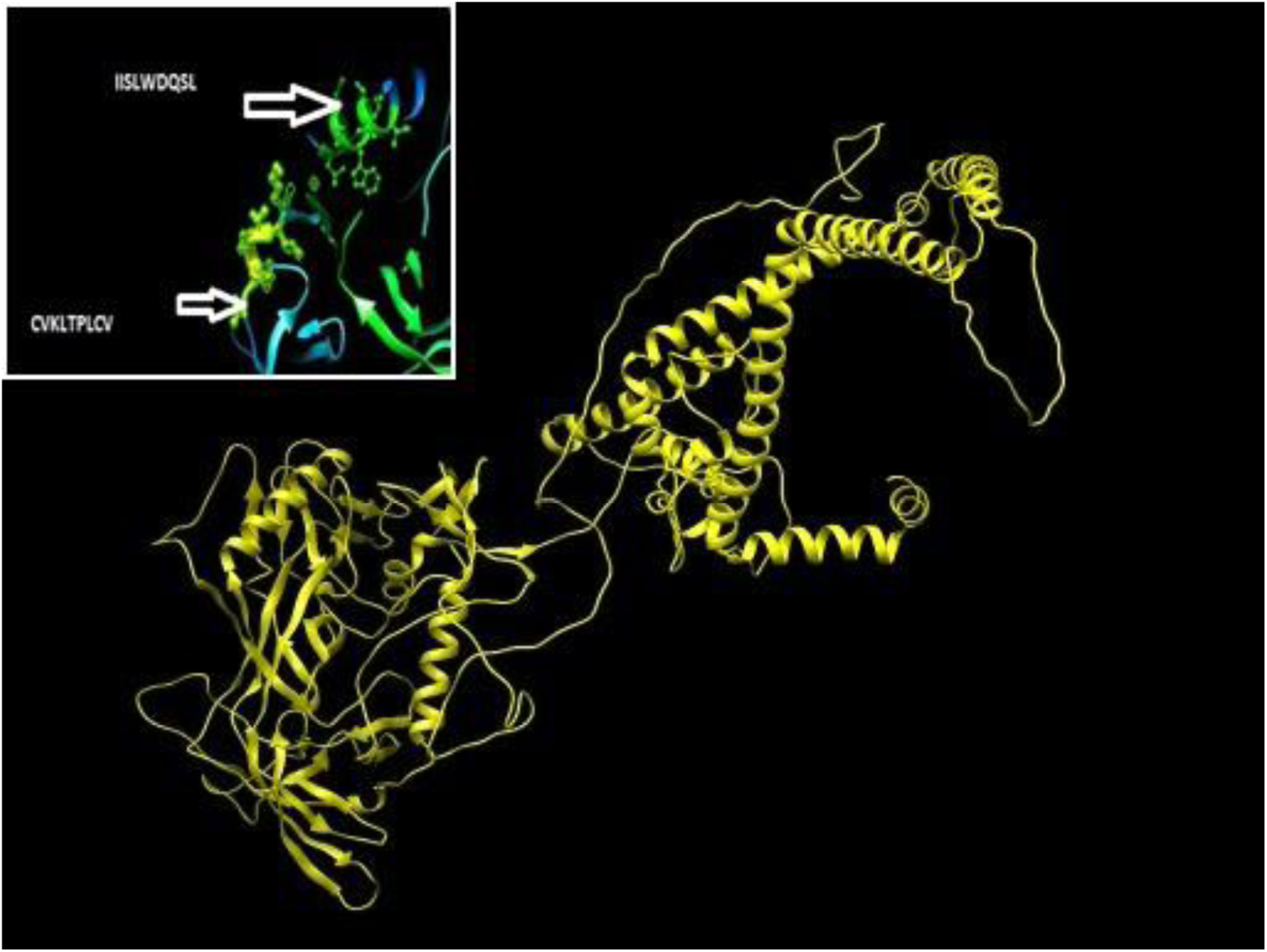
prediction peptides with MHC II of p10 by chimera 1.8

**Figure 11:**
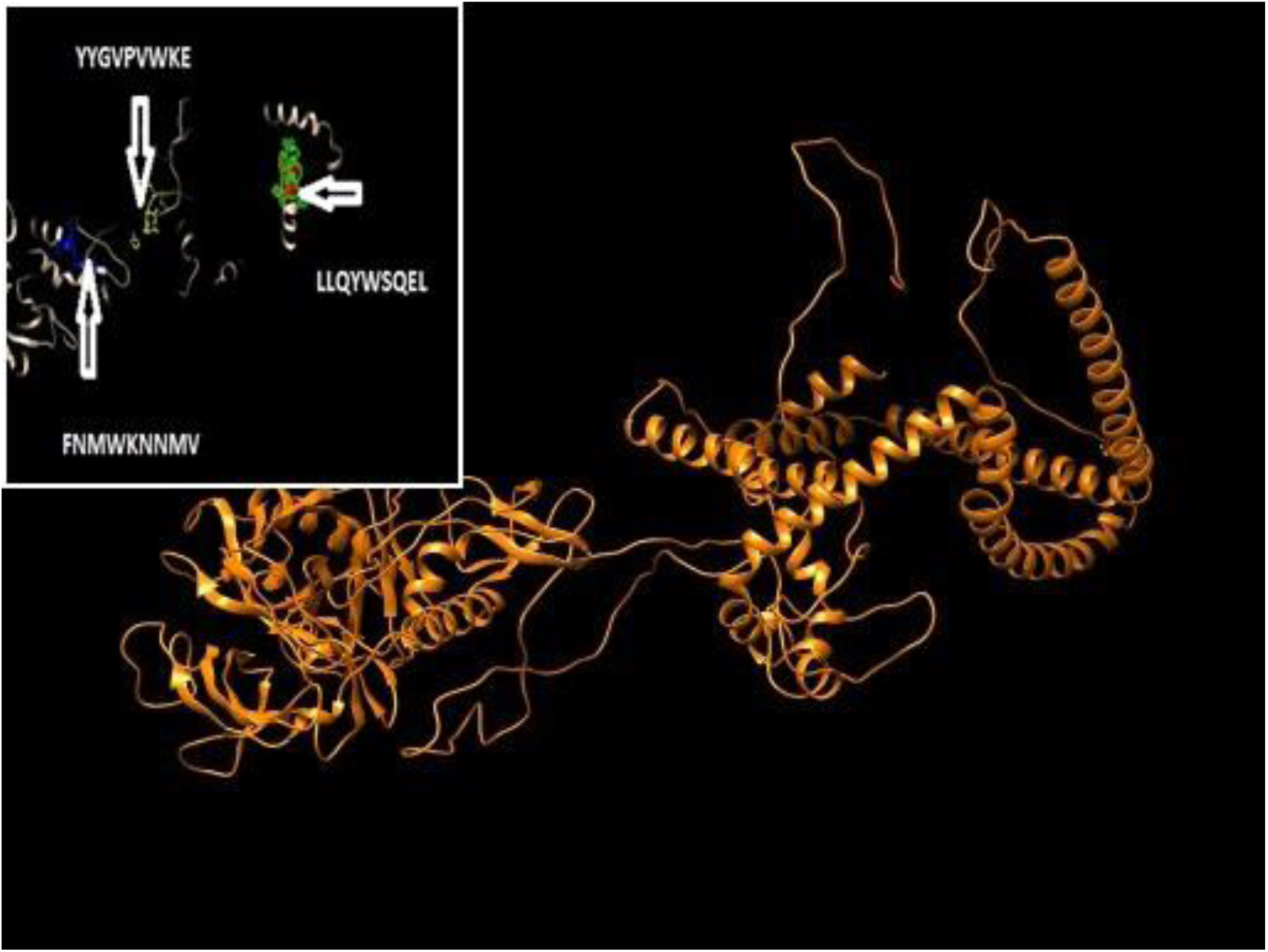
prediction peptides with MHC II of p21 by chimera 1.8

**Figure 12:**
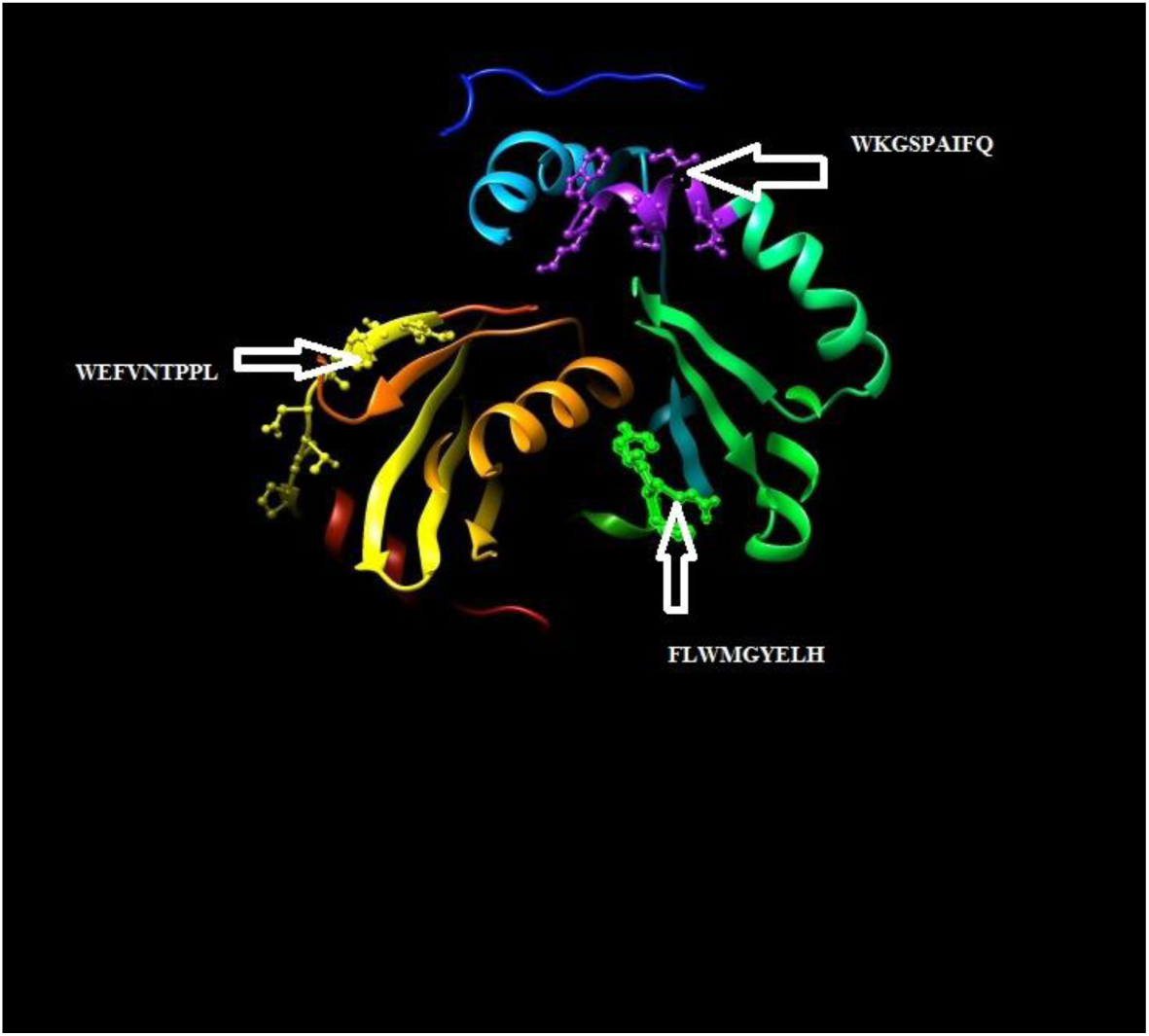
prediction peptides with MHC II of p51 by chimera 1.8

#### population coverage

Analysis of population coverage all epitopes from the reverse transcriptase proteins which selected as epitopes with high affinity to interact with MHC class I and MHC class II, subjected to IEDB population coverage set against the whole world population. For MHC class I, three Epitopes with highest population coverage resulted in epitope set of 88.87%. In MHC class II, towepitopes that interact with class II alleles. resulted in epitope set of 97.24%. for combined population coverage was also calculated for combined MHC class I and class II using proposed epitopes from reserve transcriptase protein which resulted in epitope set of 99.86%.

In p21 For MHC class I, three Epitopes with highest population coverage were. YYGVPVWKE,PQEVFLVNV and AAGSTMGAA resulted in epitope set of 97.55%. In MHC class II, three epitopes that interact with class II alleles YYGVPVWKE, FNMMWKNNMV, LLQYWSOEL resulted in epitope set of 95.94% for combined population coverage The data demonstrated in Table 7 Figure 4.3 Prediction of population coverage was also calculated for combined MHC class I and class II using proposed epitopes from structural polyprotein which resulted in epitope set of 99.62%.

In p51 For MHC class I, five Epitopes with highest population coverage resulted in epitope set of 94.45%. In MHC class II, five epitopes that interact with class II alleles. resulted in epitope set of 93.6%.for combined population coverage was also calculated for combined MHC class I and class II using proposed epitopes from reserve transcriptase protein which resulted in epitope set of 99.76%.

**Table 5:**
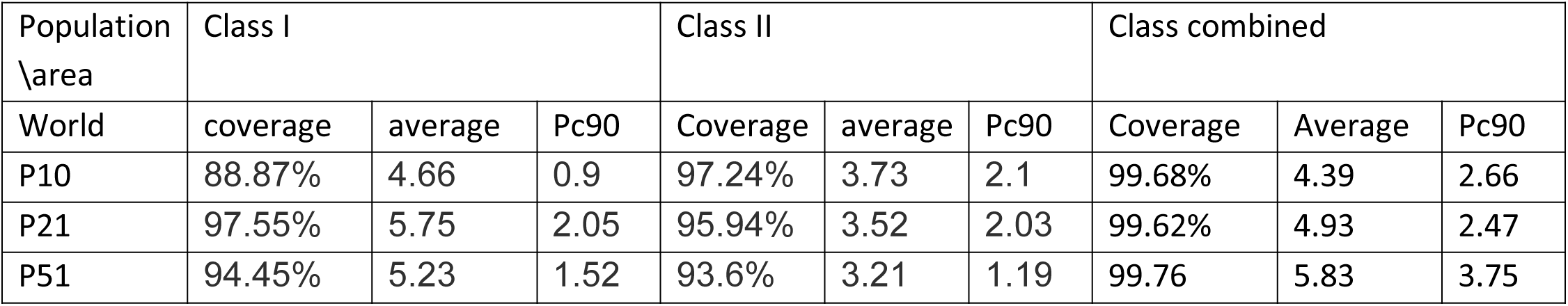
population coverage

**Figure 13:**
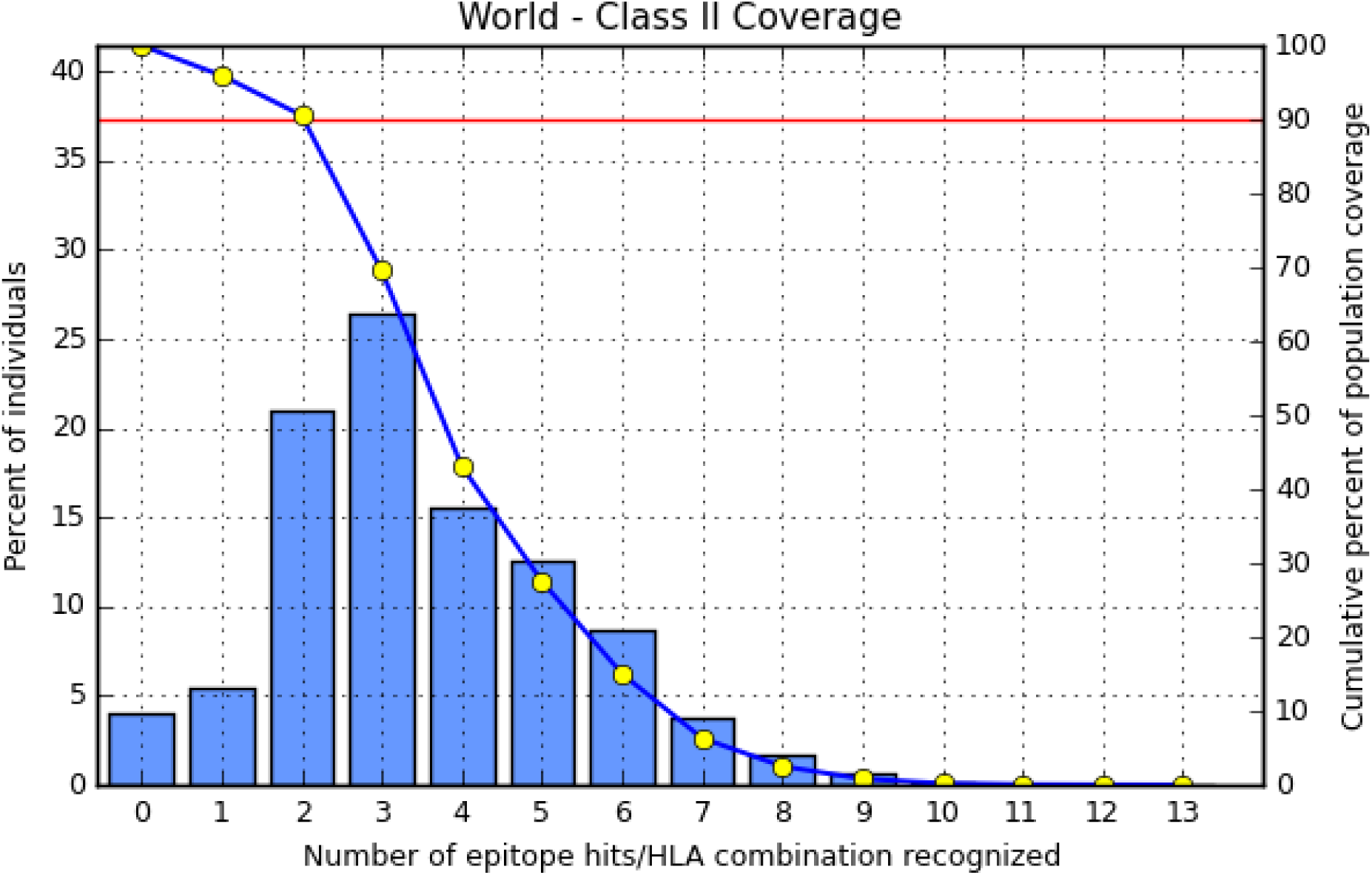
population coverage MHC class II of p10 HIV

**Figure 14:**
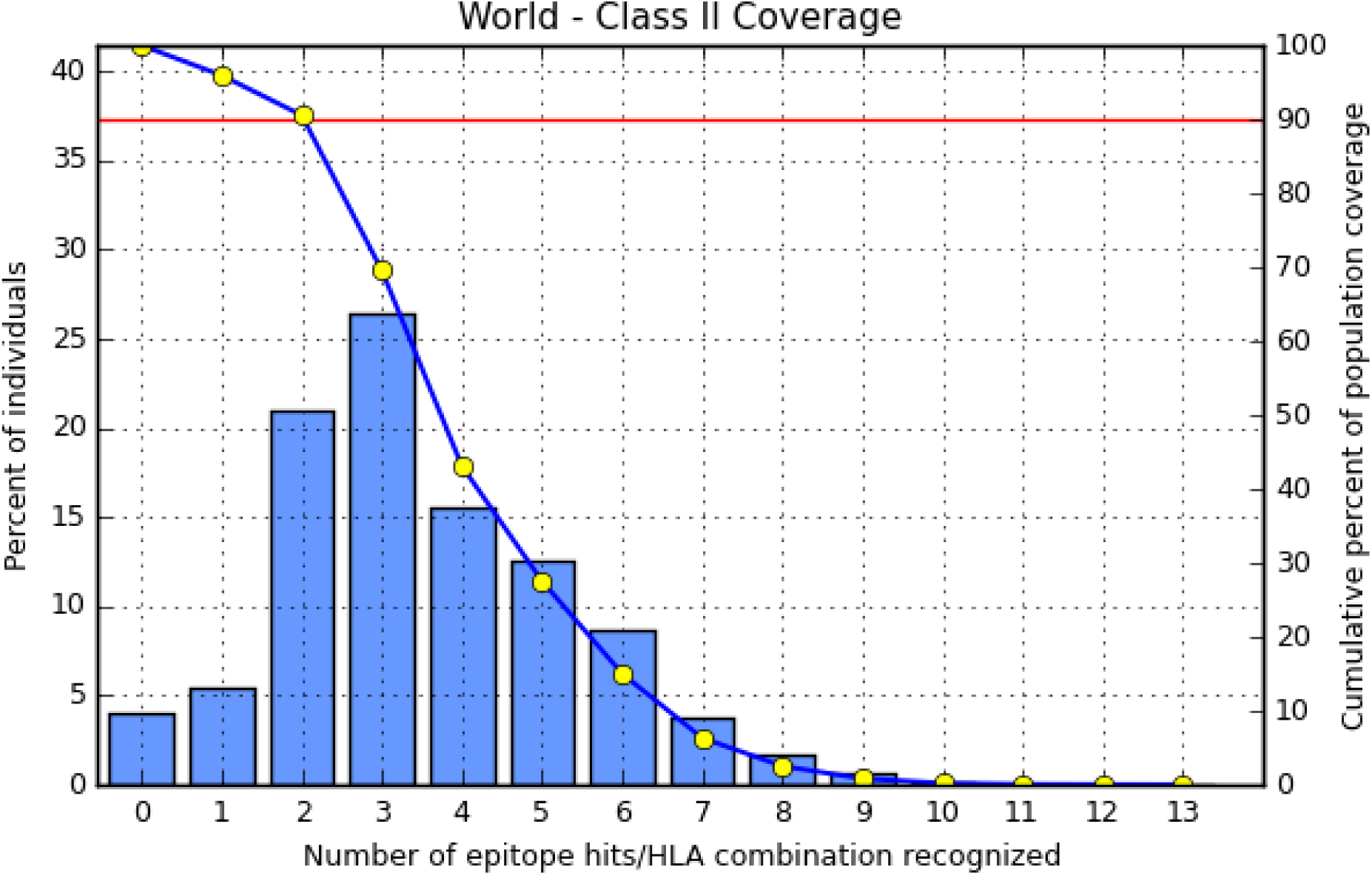
population coverage MHC class II of p21 HIV

**Figure 15:**
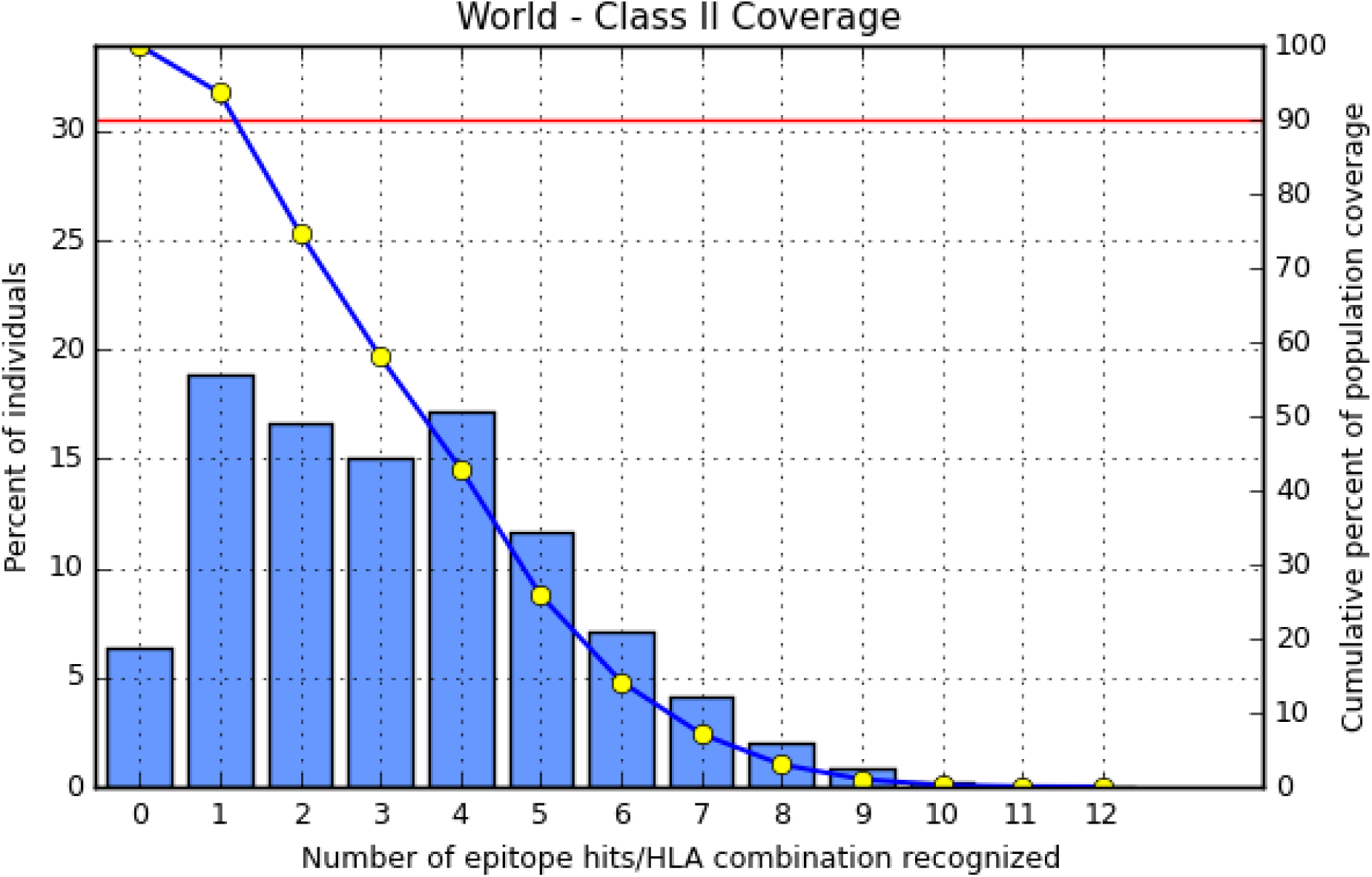
population coverage MHC class II of p51 HIV

#### Phylogenic analysis

The retrieved sequences will be conduct in Phylogenetic to determine the common ancestor of each strain using different tools from (http://www.phylogeny.fr)

**Figure 16:**
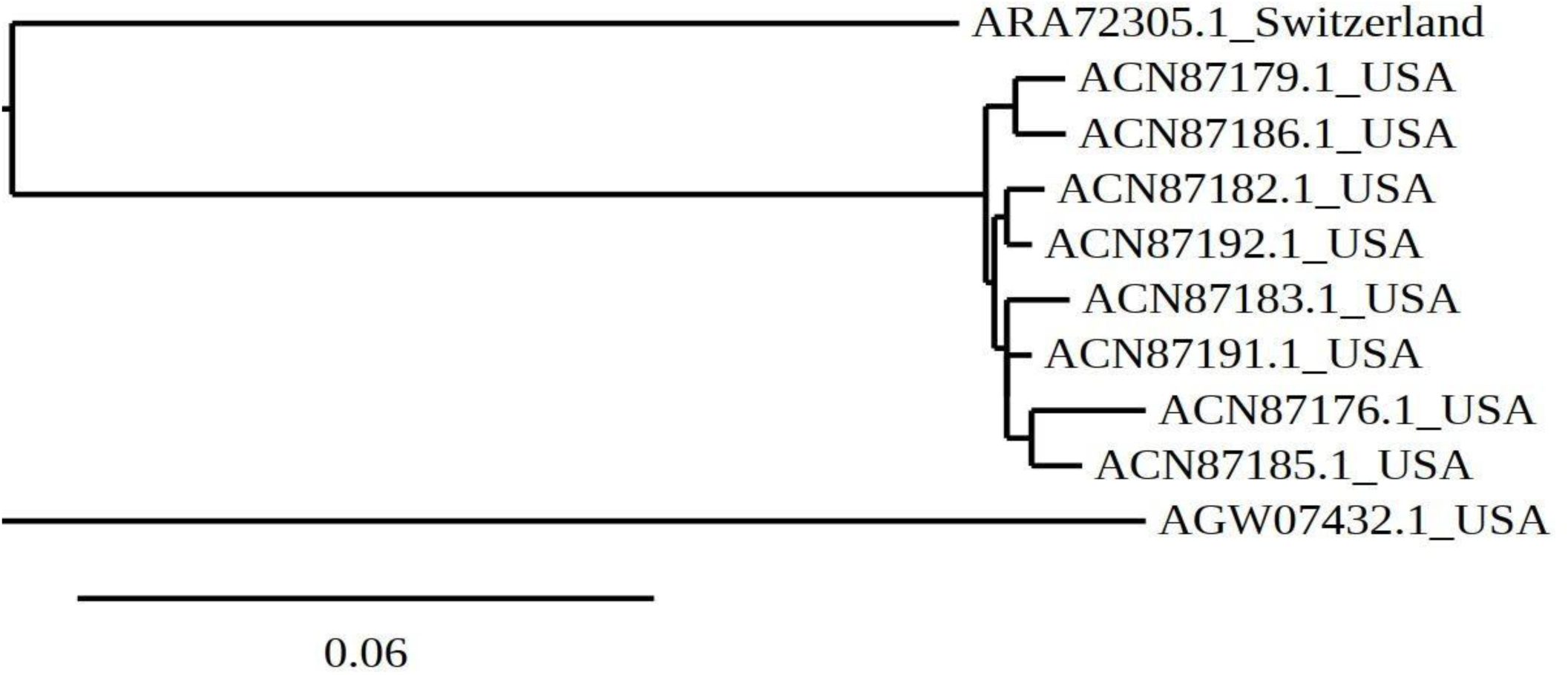
Phylogenic analysis of p10

**Figure 17:**
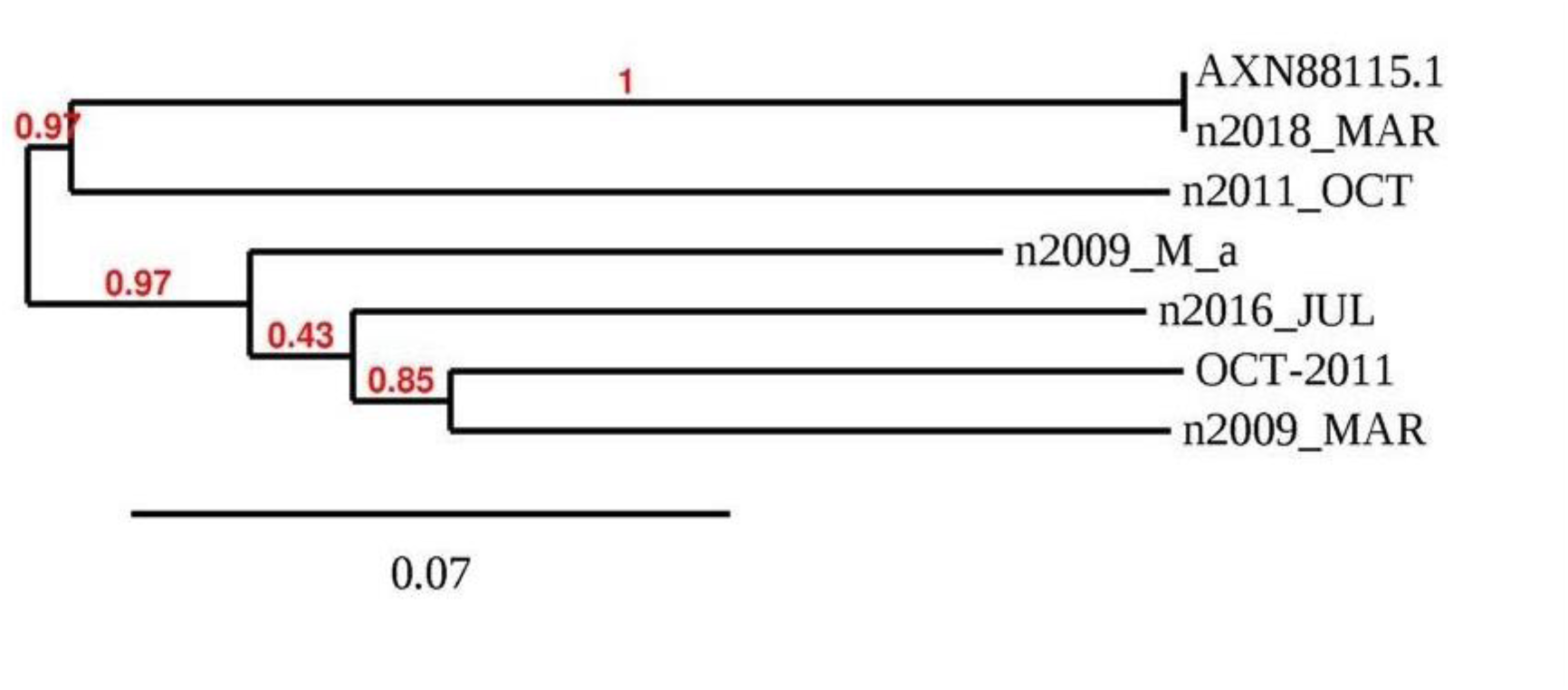
Phylogenetic aanlysis for retrieved sequences p21

**Figure 18:**
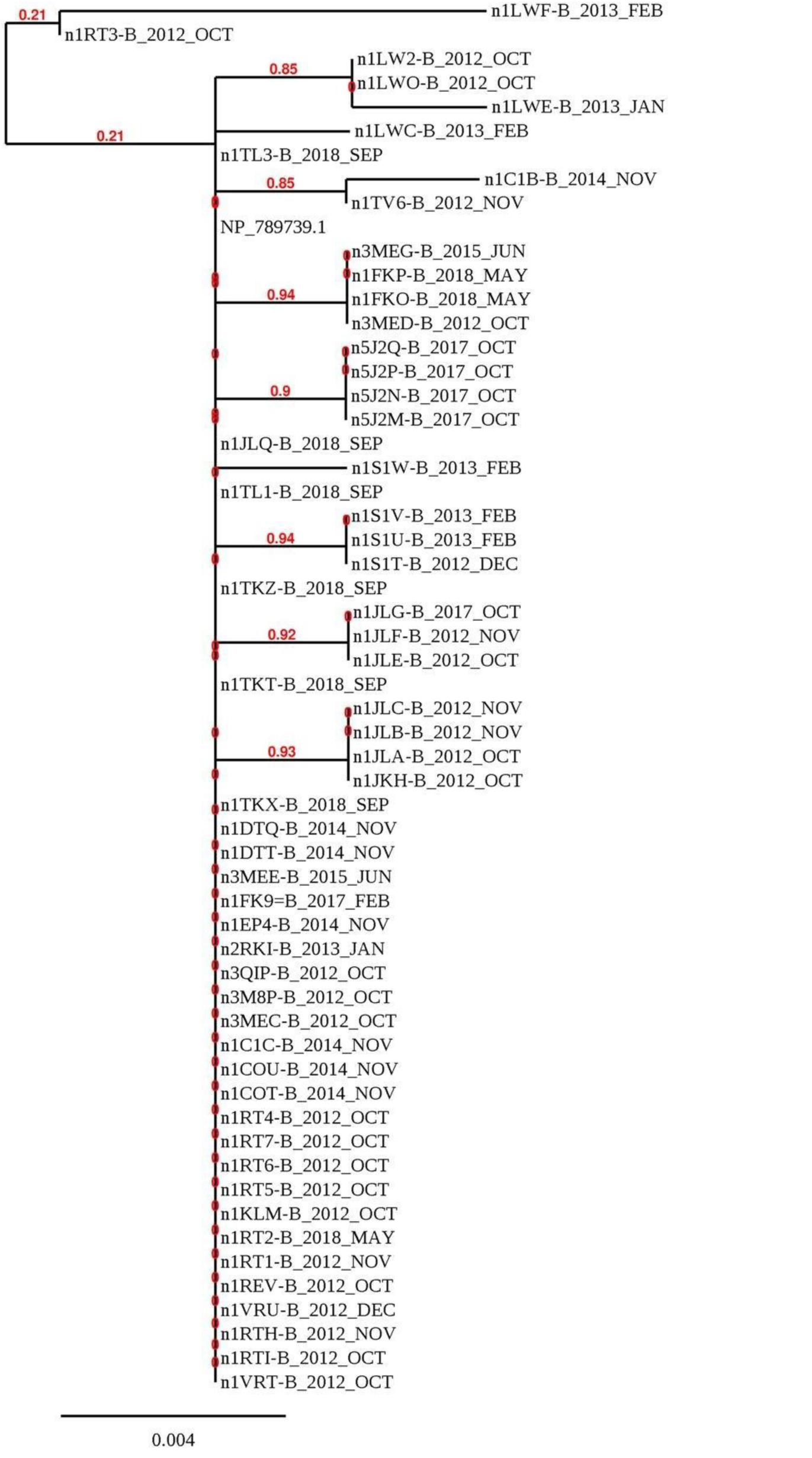
Phylogeneticaanlysis for retrieved sequence p51

## Discussion

In this study we aimed to predict possible peptide vaccine for HIV, regarding epitopes that would elicit an antibody immune response “B-cell epitopes ^708^QGYSP^712^, ^708^QGYSP^712,73^CVPTDPNPQ^81^, ^“346^**FKNL^349^**, is proposed as they got results above thresholds in Bepipred linear epitope prediction, Emini surface accessibility and Kolaskar and Tongaonkar antigenicity prediction methods and have shown 100% conservancy among the (HIV P10,P21, P51) retrieved sequences.

There were more epitopes that was addressed as potentially promising epitopes as they bound the highest number of alleles in MHC I. For p10^, 47^**EANTTLFCA**^55^, ^53^**FCASDAKAY**^61^, ^55^,**ASDAKAYE**T ^63^,from **p21** ^38^YYGVPVWKE^46^, ^10^ **PQEVFLVNV**^18^ and ^29^**AAGSTMGAA**^37^, and ^63^**`EWEFVNTPP`** ^71^epitope was found to bind 51 MHCI Alleles,^70^‘**`PPLVKLWYQ`**^78^ epitope was found tobind 50 MHC I alleles and,^79^**`LEKEPIVGA`**^87^ epitope was found to bind 51 MHCI alleles.

All epitopes were found to have a high binding affinity and lowbinding energy to MHC class I molecule in the structural level, but The^63^ **`EWEFVNTPP`^71^**, **^70^`PPLVKLWYQ`**^78^,**, ^79^`LEKEPIVGA`**^87^ epitopes were visualized because they bind the largest number of alleles, its importance as a component of the designed peptide vaccine. For the MHC II binding peptides, ^119^**IISLWDQSL***^127^,* ^108^**CVKLTPLCV***^116^for **p10***, ^38^YYGVPVWKE^46^, ^16^FNMMWKNNMV^30^,^17^LLQYWSOEL^31^,from **p21**, 174 conserved epitopes were predicted to interact with different human MHC II alleles. We postulated the best two; the 9-mer peptide (core)**^7^`WKGSPAIFQ`^21^** that showed affinity for 27 MHC- II molecules, ^11^**`FLWMGYEL`**^25^ that showed affinity for 43 MHC- II molecules,and ^58^**`WEFVNTPPL`^72^** that showed affinity for 37 MHC- II molecules.

population coverage all epitopes from the reverse transcriptase proteins which selected as epitopes with high affinity to interact with MHC class I and MHC class II, subjected to IEDB population coverage set against the whole world population. In p10 six epitopes of MHC I&MHC II was high affinity give 99.68% whole world, In p21 six epitopes of MHC I&MHC II was high affinity give 99.62% whole world,and six epitopes of MHC I&MHC II was high affinity give 99.76% whole world in p51.

**QGYSP** from 704 to 708 was found to have greatest score among all other peptides, and one peptides had got results above thresholds in Bepipred linear epitope prediction, Emini surface accessibility and Kolaskar and Tongaonkar antigenicity prediction methods and have shown 100% conservancy among the (HIV P10,P21) retrieved sequences.

The peptides found in the present study may prove more immunogenic as compared to the earlier reported peptides. Predicted peptides might show the physicochemical instability, to overcome this limitation, several structural as well as physical modification strategies are available to enhance the poor physiochemical stability of peptides. these strategies are including peptidomimetic approach, prodrug approach, analogue formations, hydrophobic ion pairing, conjugation with fatty acids and use of substitute methods of drug administration.

Researchers have been working to gather data linked to HIV to understand its biology, transmission and patho-physiology in order to eliminate the disease completely. In the near future, we anticipate that predicted epitopes have therapeutic potential with an outstanding scope.

In recent studies, novel antigenic epitopes of some essential and vital proteins revealed that can victoriously elicit response of immune system therefore becoming great peptide vaccines targets and protecting host from virus attack. So, the current research was conducted to predict antigenic epitopes of reverse transcriptaseP51 (RT) HIV. We carry out sequence, structure, and conservation analysis as well as homology modeling of reverse transcriptase P51 (RT) of HIV.

Research discloses T and B cell epitopes which are antigenic and conserved among the HIV isolates of different countries. These epitopes are capable to induce a particular immunologic response. Interestingly few of the antigenic epitopes proposed and investigated in this work might present a preliminary set of peptides for future vaccine development against HIV for control and prevention of this devastating epidemic.

## Supporting information

Sublmentary tables

