## Supplementary material for "Prediction of Multi peptides Vaccination from VP10,VP21, VP51 against Reverse Transcriptase Human immunodeficiency Viruses Using Immuno-informatics Approach": Sublmentary tables

Extra table of HIV p in MHC I

| HIV P | Allels | Start | End | Peptide | IC50 |
| --- | --- | --- | --- | --- | --- |
| P10 | HLAA*01:01 | 1 | 9 | EANTTLFCA | 14116.2 |
|  | HLAA*02:01 | 2 | 10 |  | 19889 |
|  | HLAA*02:06 | 3 | 11 |  | 7108.66 |
|  | HLAA*03:01 | 5 | 13 |  | 33202.8 |
|  | HLAA*11:01 | 6 | 14 |  | 23912.9 |
|  | HLAA*23:01 | 8 | 16 |  | 41699 |
|  | HLAA*24:02 | 9 | 17 |  | 41640.4 |
|  | HLAA*25:01 | 9 | 17 |  | 24354.4 |
|  | HLAA*26:01 | 11 | 19 |  | 18737.9 |
|  | HLAA*29:02 | 12 | 20 |  | 29021.8 |
|  | HLAA*30:01 | 13 | 21 |  | 12311.8 |
|  | HLAA*30:02 | 15 | 23 |  | 23806.8 |
|  | HLAA*31:01 | 16 | 24 |  | 29485.8 |
|  | HLAA*32:01 | 18 | 26 |  | 44399.7 |
|  | HLAA*68:01 | 18 | 26 |  | 16004.7 |
|  | HLAA*68:02 | 19 | 27 |  | 197.42 |
|  | HLAB*07:02 | 22 | 30 |  | 30713.4 |
|  | HLAB*08:01 | 22 | 30 |  | 17787.7 |
|  | HLAB*14:02 | 24 | 32 |  | 8663.05 |
|  | HLAB*15:01 | 25 | 33 |  | 27220 |
|  | HLAB*18:01 | 26 | 34 |  | 22178.5 |
|  | HLAB*27:05 | 28 | 36 |  | 28663.3 |
|  | HLAB*35:01 | 29 | 37 |  | 5546.84 |
|  | HLAB*35:03 | 30 | 38 |  | 40666.7 |
|  | HLAB*38:01 | 32 | 40 |  | 41466.9 |
|  | HLAB*39:01 | 33 | 41 |  | 18390.3 |
|  | HLAB*40:01 | 35 | 43 |  | 28579 |
|  | HLAB*40:02 | 36 | 44 |  | 36945.2 |
|  | HLAB*44:02 | 37 | 45 |  | 29349.3 |
|  | HLAB*44:03 | 38 | 46 |  | 40533.1 |
|  | HLAB*46:01 | 40 | 48 |  | 30016.3 |
|  | HLAB*48:01 | 41 | 49 |  | 37808.6 |
|  | HLAB*51:01 | 42 | 50 |  | 21010.1 |
|  | HLAB*53:01 | 43 | 51 |  | 11302.6 |
|  | HLAB*57:01 | 45 | 53 |  | 24916.3 |
|  | HLAB*58:01 | 46 | 54 |  | 9983.09 |
|  | HLAB*58:02 | 48 | 56 |  | 37206 |
|  | HLAC*03:03 | 48 | 56 |  | 24655.6 |
|  | HLAC*04:01 | 50 | 58 |  | 28242.4 |
|  | HLAC*05:01 | 51 | 59 |  | 21701.6 |
|  | HLAC*06:02 | 53 | 61 |  | 34218.6 |
|  | HLAC*07:01 | 54 | 62 |  | 32010.2 |
|  | HLAC*07:02 | 55 | 63 |  | 36434.8 |
|  | HLAC*08:02 | 56 | 64 |  | 33176.3 |
|  | HLAC*12:03 | 58 | 66 |  | 31279.2 |
|  | HLAC*14:02 | 60 | 68 |  | 40687.4 |
|  | HLAC*15:02 | 60 | 68 |  | 13335.7 |
|  | HLAE*01:01 | 62 | 70 |  | 42088 |
|  | HLAA*01:01 | 1 | 9 | FCASDAKAY | 13382.5 |
|  | HLAA*02:01 | 2 | 10 |  | 33328.5 |
|  | HLAA*02:06 | 4 | 12 |  | 36915.7 |
|  | HLAA*03:01 | 5 | 13 |  | 31878.5 |
|  | HLAA*11:01 | 6 | 14 |  | 34887.4 |
|  | HLAA*23:01 | 7 | 15 |  | 39951.4 |
|  | HLAA*24:02 | 9 | 17 |  | 42691.1 |
|  | HLAA*25:01 | 9 | 17 |  | 8405.08 |
|  | HLAA*26:01 | 11 | 19 |  | 16738.5 |
|  | HLAA*29:02 | 12 | 20 |  | 1061.67 |
|  | HLAA*30:01 | 14 | 22 |  | 31969.7 |
|  | HLAA*30:02 | 14 | 22 |  | 621.64 |
|  | HLAA*31:01 | 17 | 25 |  | 40067.9 |
|  | HLAA*32:01 | 18 | 26 |  | 41082.8 |
|  | HLAA*68:01 | 18 | 26 |  | 23272.2 |
|  | HLAA*68:02 | 20 | 28 |  | 36302.5 |
|  | HLAB*07:02 | 21 | 29 |  | 30310.7 |
|  | HLAB*08:01 | 23 | 31 |  | 29307.1 |
|  | HLAB*14:02 | 24 | 32 |  | 13455.2 |
|  | HLAB*15:01 | 25 | 33 |  | 3406.66 |
|  | HLAB*18:01 | 26 | 34 |  | 9954.94 |
|  | HLAB*27:05 | 28 | 36 |  | 22300.7 |
|  | HLAB*35:01 | 29 | 37 |  | 13.55 |
|  | HLAB*35:03 | 30 | 38 |  | 30924.9 |
|  | HLAB*38:01 | 32 | 40 |  | 31561.7 |
|  | HLAB*39:01 | 33 | 41 |  | 21489.4 |
|  | HLAB*40:01 | 34 | 42 |  | 21132 |
|  | HLAB*40:02 | 36 | 44 |  | 44852.6 |
|  | HLAB*44:02 | 36 | 44 |  | 9349.02 |
|  | HLAB*44:03 | 38 | 46 |  | 23936.2 |
|  | HLAB*46:01 | 39 | 47 |  | 11229.7 |
|  | HLAB*48:01 | 41 | 49 |  | 37237.4 |
|  | HLAB*51:01 | 42 | 50 |  | 27206.2 |
|  | HLAB*53:01 | 43 | 51 |  | 5480.26 |
|  | HLAB*57:01 | 45 | 53 |  | 20197.6 |
|  | HLAB*58:01 | 46 | 54 |  | 23334.5 |
|  | HLAB*58:02 | 47 | 55 |  | 32964.1 |
|  | HLAC*03:03 | 48 | 56 |  | 327.72 |
|  | HLAC*04:01 | 50 | 58 |  | 19572 |
| P21 | HLA-A*01:01 | 10 | 18 | PQEVFLVNV | 22120.48 |
|  | HLA-A*02:01 | 10 | 18 |  | 12344.89 |
|  | HLA-A*02:06 | 10 | 18 |  | 1220.83 |
|  | HLA-A*03:01 | 10 | 18 |  | 33440.43 |
|  | HLA-A*11:01 | 10 | 18 |  | 34849.64 |
|  | HLA-A*23:01 | 10 | 18 |  | 28660.15 |
|  | HLA-A*24:02 | 10 | 18 |  | 39171.12 |
|  | HLA-A*25:01 | 10 | 18 |  | 36582.49 |
|  | HLA-A*26:01 | 10 | 18 |  | 30930.53 |
|  | HLA-A*29:02 | 10 | 18 |  | 28242.73 |
|  | HLA-A*30:01 | 10 | 18 |  | 23754.54 |
|  | HLA-A*30:02 | 10 | 18 |  | 26213.85 |
|  | HLA-A*31:01 | 10 | 18 |  | 26915.7 |
|  | HLA-A*32:01 | 10 | 18 |  | 29995.85 |
|  | HLA-A*68:01 | 10 | 18 |  | 44333.92 |
|  | HLA-A*68:02 | 10 | 18 |  | 31054.61 |
|  | HLA-B*07:02 | 10 | 18 |  | 36332 |
|  | HLA-B*08:01 | 10 | 18 |  | 32236.77 |
|  | HLA-B*14:02 | 10 | 18 |  | 28903.37 |
|  | HLA-B*15:01 | 10 | 18 |  | 29934.57 |
|  | HLA-B*15:02 | 10 | 18 |  | 37507.93 |
|  | HLA-B*15:02 | 10 | 18 |  | 37507.93 |
|  | HLA-B*18:01 | 10 | 18 |  | 30154.97 |
|  | HLA-B*27:05 | 10 | 18 |  | 33989.4 |
|  | HLA-B*35:01 | 10 | 18 |  | 42110.72 |
|  | HLA-B*35:03 | 10 | 18 |  | 44129.11 |
|  | HLA-B*39:01 | 10 | 18 |  | 27324.15 |
|  | HLA-B*40:01 | 10 | 18 |  | 16159.77 |
|  | HLA-B*40:02 | 10 | 18 |  | 28201.51 |
|  | HLA-B*44:02 | 10 | 18 |  | 38075.47 |
|  | HLA-B*44:03 | 10 | 18 |  | 41428.3 |
|  | HLA-B*46:01 | 10 | 18 |  | 32761.04 |
|  | HLA-B*48:01 | 10 | 18 |  | 25356.04 |
|  | HLA-B*51:01 | 10 | 18 |  | 28624.5 |
|  | HLA-B*53:01 | 10 | 18 |  | 39960.93 |
|  | HLA-B*57:01 | 10 | 18 |  | 32840.2 |
|  | HLA-B*58:01 | 10 | 18 |  | 30820.3 |
|  | HLA-B*58:02 | 10 | 18 |  | 38078.76 |
|  | HLA-C*03:03 | 10 | 18 |  | 46974.44 |
|  | HLA-C*04:01 | 10 | 18 |  | 30678.89 |
|  | HLA-C*05:01 | 10 | 18 |  | 42626.43 |
|  | HLA-C*06:02 | 10 | 18 |  | 40444.62 |
|  | HLA-C*07:01 | 10 | 18 |  | 35748.64 |
|  | HLA-C*07:02 | 10 | 18 |  | 40403.51 |
|  | HLA-C*08:02 | 10 | 18 |  | 34868.12 |
|  | HLA-C*12:03 | 10 | 18 |  | 18205.15 |
|  | HLA-C*14:02 | 10 | 18 |  | 42184.6 |
|  | HLA-C*15:02 | 10 | 18 |  | 28389.79 |
|  | HLA-E*01:01 | 10 | 18 |  | 44402.11 |
|  | HLA-A*01:01 | 38 | 46 | YYGVPVWKE | 30072.22 |
|  | HLA-A*02:01 | 38 | 46 |  | 32155.27 |
|  | HLA-A*02:06 | 38 | 46 |  | 24646.01 |
|  | HLA-A*03:01 | 38 | 46 |  | 39504.37 |
|  | HLA-A*11:01 | 38 | 46 |  | 33742.09 |
|  | HLA-A*23:01 | 38 | 46 |  | 2526.83 |
|  | HLA-A*24:02 | 38 | 46 |  | 5342.85 |
|  | HLA-A*25:01 | 38 | 46 |  | 32689.16 |
|  | HLA-A*26:01 | 38 | 46 |  | 34788.61 |
|  | HLA-A*29:02 | 38 | 46 |  | 10098.58 |
|  | HLA-A*30:01 | 38 | 46 |  | 31227.45 |
|  | HLA-A*30:02 | 38 | 46 |  | 21239.55 |
|  | HLA-A*31:01 | 38 | 46 |  | 22639.37 |
|  | HLA-A*32:01 | 38 | 46 |  | 40533.12 |
|  | HLA-A*68:01 | 38 | 46 |  | 36913.28 |
|  | HLA-A*68:02 | 38 | 46 |  | 33317.62 |
|  | HLA-B*07:02 | 38 | 46 |  | 37750.57 |
|  | HLA-B*08:01 | 38 | 46 |  | 31149.51 |
|  | HLA-B*14:02 | 38 | 46 |  | 29143.9 |
|  | HLA-B*15:01 | 38 | 46 |  | 38305.21 |
|  | HLA-B*15:02 | 38 | 46 |  | 29538.83 |
|  | HLA-B*15:02 | 38 | 46 |  | 29538.83 |
|  | HLA-B*18:01 | 38 | 46 |  | 23571.49 |
|  | HLA-B*27:05 | 38 | 46 |  | 33366.32 |
|  | HLA-B*35:01 | 38 | 46 |  | 37062.56 |
|  | HLA-B*35:03 | 38 | 46 |  | 44969.7 |
|  | HLA-B*39:01 | 38 | 46 |  | 34226.7 |
|  | HLA-B*40:01 | 38 | 46 |  | 25634.93 |
|  | HLA-B*40:02 | 38 | 46 |  | 41976.07 |
|  | HLA-B*44:02 | 38 | 46 |  | 33489.31 |
|  | HLA-B*44:03 | 38 | 46 |  | 45379.32 |
|  | HLA-B*46:01 | 38 | 46 |  | 31945.45 |
|  | HLA-B*48:01 | 38 | 46 |  | 34530.6 |
|  | HLA-B*51:01 | 38 | 46 |  | 34539.2 |
|  | HLA-B*53:01 | 38 | 46 |  | 34275.63 |
|  | HLA-B*57:01 | 38 | 46 |  | 30950.62 |
|  | HLA-B*58:01 | 38 | 46 |  | 25579.79 |
|  | HLA-B*58:02 | 38 | 46 |  | 36731.21 |
|  | HLA-C*03:03 | 38 | 46 |  | 35619.31 |
|  | HLA-C*04:01 | 38 | 46 |  | 24359.16 |
|  | HLA-C*05:01 | 38 | 46 |  | 41621.06 |
|  | HLA-C*06:02 | 38 | 46 |  | 16337.51 |
|  | HLA-C*07:01 | 38 | 46 |  | 14278.38 |
|  | HLA-C*07:02 | 38 | 46 |  | 3555.02 |
|  | HLA-C*08:02 | 38 | 46 |  | 38764.26 |
|  | HLA-C*12:03 | 38 | 46 |  | 26367.46 |
|  | HLA-C*14:02 | 38 | 46 |  | 7256.47 |
|  | HLA-C*15:02 | 38 | 46 |  | 41203.9 |
|  | HLA-E*01:01 | 38 | 46 |  | 44182.13 |
| P51 | HLA-A*01:01 | 71 | 63 | EWEFVNTPP | 25684.62 |
|  | HLA-A*02:01 | 71 | 63 |  | 40081.3 |
|  | HLA-A*02:06 | 71 | 63 |  | 36641.5 |
|  | HLA-A*03:01 | 71 | 63 |  | 43600.4 |
|  | HLA-A*11:01 | 71 | 63 |  | 43301.4 |
|  | HLA-A*23:01 | 71 | 63 |  | 30989.2 |
|  | HLA-A*24:02 | 71 | 63 |  | 31579.8 |
|  | HLA-A*25:01 | 71 | 63 |  | 44826.9 |
|  | HLA-A*26:01 | 71 | 63 |  | 43630.1 |
|  | HLA-A*29:02 | 71 | 63 |  | 37443.5 |
|  | HLA-A*30:01 | 71 | 63 |  | 24782.7 |
|  | HLA-A*30:02 | 71 | 63 |  | 35247.8 |
|  | HLA-A*31:01 | 71 | 63 |  | 36081.7 |
|  | HLA-A*32:01 | 71 | 63 |  | 44275.9 |
|  | HLA-A*33:03 | 71 | 63 |  | 30109 |
|  | HLA-A*68:01 | 71 | 63 |  | 35734.4 |
|  | HLA-A*68:02 | 71 | 63 |  | 27566.5 |
|  | HLA-A*74:01 | 71 | 63 |  | 43521.2 |
|  | HLA-B*07:02 | 71 | 63 |  | 42969.1 |
|  | HLA-B*08:01 | 71 | 63 |  | 30663.95 |
|  | HLA-B*14:02 | 71 | 63 |  | 17055.49 |
|  | HLA-B*15:01 | 71 | 63 |  | 30725.06 |
|  | HLA-B*15:02 | 71 | 63 |  | 30594.36 |
|  | HLA-B*18:01 | 71 | 63 |  | 23130.86 |
|  | HLA-B*27:05 | 71 | 63 |  | 28980.39 |
|  | HLA-B*35:01 | 71 | 63 |  | 19464.29 |
|  | HLA-B*35:03 | 71 | 63 |  | 44218.96 |
|  | HLA-B*38:01 | 71 | 63 |  | 44950.74 |
|  | HLA-B*39:01 | 71 | 63 |  | 27632.76 |
|  | HLA-B*40:01 | 71 | 63 |  | 19984.79 |
|  | HLA-B*40:02 | 71 | 63 |  | 45053.97 |
|  | HLA-B*44:02 | 71 | 63 |  | 29755.36 |
|  | HLA-B*44:03 | 71 | 63 |  | 44296.05 |
|  | HLA-B*48:01 | 71 | 63 |  | 35418.31 |
|  | HLA-B*46:01 | 71 | 63 |  | 32821.01 |
|  | HLA-B*51:01 | 71 | 63 |  | 31641.71 |
|  | HLA-B*53:01 | 71 | 63 |  | 28360.02 |
|  | HLA-B*57:01 | 71 | 63 |  | 35089.51 |
|  | HLA-B*58:01 | 71 | 63 |  | 31987.3 |
|  | HLA-B*58:02 | 71 | 63 |  | 37358.08 |
|  | HLA-C*03:03 | 71 | 63 |  | 34791.23 |
|  | HLA-C*04:01 | 71 | 63 |  | 16714.96 |
|  | HLA-C*05:01 | 71 | 63 |  | 36992.85 |
|  | HLA-C*06:02 | 71 | 63 |  | 47621.79 |
|  | HLA-C*07:01 | 71 | 63 |  | 45514.05 |
|  | HLA-C*07:02 | 71 | 63 |  | 40462.57 |
|  | HLA-C*08:02 | 71 | 63 |  | 40144.25 |
|  | HLA-C*12:03 | 71 | 63 |  | 36892.93 |
|  | HLA-C*14:02 | 71 | 63 |  | 27258.89 |
|  | HLA-C*15:02 | 71 | 63 |  | 42899.43 |
|  | HLA-E*01:01 | 71 | 63 |  | 44677.74 |
|  | HLA-A*01:01 | 87 | 79 | LEKEPIVGA | 33982.43 |
|  | HLA-A*02:01 | 87 | 79 |  | 29662.1 |
|  | HLA-A*02:06 | 87 | 79 |  | 14129.8 |
|  | HLA-A*03:01 | 87 | 79 |  | 38603.1 |
|  | HLA-A*11:01 | 87 | 79 |  | 40607.3 |
|  | HLA-A*23:01 | 87 | 79 |  | 43799 |
|  | HLA-A*24:02 | 87 | 79 |  | 44948.8 |
|  | HLA-A*25:01 | 87 | 79 |  | 39340.6 |
|  | HLA-A*26:01 | 87 | 79 |  | 35894.8 |
|  | HLA-A*29:02 | 87 | 79 |  | 40359.4 |
|  | HLA-A*30:01 | 87 | 79 |  | 16503.6 |
|  | HLA-A*30:02 | 87 | 79 |  | 30716.1 |
|  | HLA-A*31:01 | 87 | 79 |  | 35843.5 |
|  | HLA-A*32:01 | 87 | 79 |  | 42515 |
|  | HLA-A*33:03 | 87 | 79 |  | 38966.9 |
|  | HLA-A*68:01 | 87 | 79 |  | 39416.9 |
|  | HLA-A*68:02 | 87 | 79 |  | 24374.2 |
|  | HLA-A*74:01 | 87 | 79 |  | 42731.3 |
|  | HLA-B*07:02 | 87 | 79 |  | 31620.1 |
|  | HLA-B*08:01 | 87 | 79 |  | 26544.06 |
|  | HLA-B*14:02 | 87 | 79 |  | 24853.27 |
|  | HLA-B*15:01 | 87 | 79 |  | 11627.66 |
|  | HLA-B*15:02 | 87 | 79 |  | 35030.34 |
|  | HLA-B*18:01 | 87 | 79 |  | 6893.25 |
|  | HLA-B*27:05 | 87 | 79 |  | 29086.9 |
|  | HLA-B*35:01 | 87 | 79 |  | 40460.38 |
|  | HLA-B*35:03 | 87 | 79 |  | 45985.77 |
|  | HLA-B*38:01 | 87 | 79 |  | 45432.38 |
|  | HLA-B*39:01 | 87 | 79 |  | 37611.55 |
|  | HLA-B*40:01 | 87 | 79 |  | 11041.71 |
|  | HLA-B*40:02 | 87 | 79 |  | 4002.35 |
|  | HLA-B*44:02 | 87 | 79 |  | 11425.38 |
|  | HLA-B*44:03 | 87 | 79 |  | 27710.3 |
|  | HLA-B*48:01 | 87 | 79 |  | 28290.75 |
|  | HLA-B*46:01 | 87 | 79 |  | 30676.57 |
|  | HLA-B*51:01 | 87 | 79 |  | 33342.51 |
|  | HLA-B*53:01 | 87 | 79 |  | 43596.15 |
|  | HLA-B*57:01 | 87 | 79 |  | 31606.81 |
|  | HLA-B*58:01 | 87 | 79 |  | 31801.65 |
|  | HLA-B*58:02 | 87 | 79 |  | 28364 |
|  | HLA-C*03:03 | 87 | 79 |  | 42081.11 |
|  | HLA-C*04:01 | 87 | 79 |  | 19791.55 |
|  | HLA-C*05:01 | 87 | 79 |  | 36814.36 |
|  | HLA-C*06:02 | 87 | 79 |  | 42941.66 |
|  | HLA-C*07:01 | 87 | 79 |  | 37103.88 |
|  | HLA-C*07:02 | 87 | 79 |  | 38012.49 |
|  | HLA-C*08:02 | 87 | 79 |  | 43534.4 |
|  | HLA-C*12:03 | 87 | 79 |  | 15402.15 |
|  | HLA-C*14:02 | 87 | 79 |  | 45628.43 |
|  | HLA-C*15:02 | 87 | 79 |  | 46522.75 |
|  | HLA-E*01:01 | 87 | 79 |  | 44573.47 |

Exstra table of HIV p in MHC II

| HIV p | Allels | Start | End | Core | Peptide | IC50 |
| --- | --- | --- | --- | --- | --- | --- |
| P10 | HLADPA1*03:01/DPB1*04:02 | 35 | 49 | IISLWDQSL | HEDIISLWDQSLKPC | 331.6 |
|  | HLADPA1*03:01/DPB1*04:02 | 34 | 48 |  | MHEDIISLWDQSLKP | 380.8 |
|  | HLADQA1*01:01/DQB1*05:01 | 36 | 50 |  | EDIISLWDQSLKPCV | 185.2 |
|  | HLADQA1*01:01/DQB1*05:01 | 35 | 49 |  | HEDIISLWDQSLKPC | 186.6 |
|  | HLADQA1*01:01/DQB1*05:01 | 34 | 48 |  | MHEDIISLWDQSLKP | 191.4 |
|  | HLADQA1*01:01/DQB1*05:01 | 33 | 47 |  | QMHEDIISLWDQSLK | 234.3 |
|  | HLADQA1*01:01/DQB1*05:01 | 37 | 51 |  | DIISLWDQSLKPCVK | 252.5 |
|  | HLADQA1*01:01/DQB1*05:01 | 32 | 46 |  | DQMHEDIISLWDQSL | 274.6 |
|  | HLADQA1*05:01/DQB1*02:01 | 32 | 46 |  | DQMHEDIISLWDQSL | 456.4 |
|  | HLA-DRB1*01:01 | 35 | 49 |  | HEDIISLWDQSLKPC | 72 |
|  | HLA-DRB1*01:01 | 33 | 47 |  | QMHEDIISLWDQSLK | 97.9 |
|  | HLA-DRB1*01:01 | 34 | 48 |  | MHEDIISLWDQSLKP | 114.8 |
|  | HLA-DRB1*01:01 | 36 | 50 |  | EDIISLWDQSLKPCV | 163.4 |
|  | HLA-DRB1*01:01 | 32 | 46 |  | DQMHEDIISLWDQSL | 266.1 |
|  | HLA-DRB1*01:01 | 38 | 52 |  | IISLWDQSLKPCVKL | 304.1 |
|  | HLA-DRB1*01:01 | 37 | 51 |  | DIISLWDQSLKPCVK | 314.1 |
|  | HLA-DRB1*04:05 | 33 | 47 |  | QMHEDIISLWDQSLK | 273.1 |
|  | HLA-DRB1*04:05 | 32 | 46 |  | DQMHEDIISLWDQSL | 298.3 |
|  | HLA-DRB1*04:05 | 34 | 48 |  | MHEDIISLWDQSLKP | 361.6 |
|  | HLA-DRB1*04:05 | 35 | 49 |  | HEDIISLWDQSLKPC | 391.4 |
|  | HLA-DRB1*04:05 | 36 | 50 |  | EDIISLWDQSLKPCV | 466.9 |
|  | HLA-DRB1*15:01 | 36 | 50 |  | EDIISLWDQSLKPCV | 206.8 |
|  | HLA-DRB1*15:01 | 38 | 52 |  | IISLWDQSLKPCVKL | 230.9 |
|  | HLA-DRB1*15:01 | 37 | 51 |  | DIISLWDQSLKPCVK | 235.1 |
|  | HLA-DRB1*15:01 | 34 | 48 |  | MHEDIISLWDQSLKP | 241.1 |
|  | HLA-DRB1*15:01 | 35 | 49 |  | HEDIISLWDQSLKPC | 242.4 |
|  | HLA-DRB1*15:01 | 33 | 47 |  | QMHEDIISLWDQSLK | 286.7 |
|  | HLA-DRB4*01:01 | 35 | 49 |  | HEDIISLWDQSLKPC | 125.8 |
|  | HLA-DRB4*01:01 | 36 | 50 |  | EDIISLWDQSLKPCV | 127.1 |
|  | HLA-DRB4*01:01 | 34 | 48 |  | MHEDIISLWDQSLKP | 132.8 |
|  | HLA-DRB4*01:01 | 33 | 47 |  | QMHEDIISLWDQSLK | 154.3 |
|  | HLA-DRB4*01:01 | 37 | 51 |  | DIISLWDQSLKPCVK | 183.7 |
|  | HLA-DRB4*01:01 | 32 | 46 |  | DQMHEDIISLWDQSL | 206.1 |
|  | HLA-DRB4*01:01 | 38 | 52 |  | IISLWDQSLKPCVKL | 255.9 |
| P21 | HLA-DPA1*01/DPB1*04:01 | 32 | 46 | YYGVPVWKE | NWWVTVYYGVPVWKE | 61.8 |
|  | HLA-DPA1*01:03/DPB1*02:01 | 32 | 46 |  | NWWVTVYYGVPVWKE | 42.4 |
|  | HLA-DPA1*02:01/DPB1*01:01 | 32 | 46 |  | NWWVTVYYGVPVWKE | 103.6 |
|  | HLA-DRB1*04:01 | 32 | 46 |  | NWWVTVYYGVPVWKE | 397.7 |
|  | HLA-DRB1*04:05 | 32 | 46 |  | NWWVTVYYGVPVWKE | 248.1 |
|  | HLA-DRB1*07:01 | 32 | 46 |  | NWWVTVYYGVPVWKE | 312 |
|  | HLA-DPA1*01/DPB1*04:01 | 33 | 47 |  | WWVTVYYGVPVWKEA | 51.7 |
|  | HLA-DPA1*01:03/DPB1*02:01 | 33 | 47 |  | WWVTVYYGVPVWKEA | 50.5 |
|  | HLA-DPA1*02:01/DPB1*01:01 | 33 | 47 |  | WWVTVYYGVPVWKEA | 87.2 |
|  | HLA-DQA1*04:01/DQB1*04:02 | 33 | 47 |  | WWVTVYYGVPVWKEA | 413.1 |
|  | HLA-DRB1*01:01 | 33 | 47 |  | WWVTVYYGVPVWKEA | 116.1 |
|  | HLA-DRB1*04:01 | 33 | 47 |  | WWVTVYYGVPVWKEA | 285.1 |
|  | HLA-DRB1*04:05 | 33 | 47 |  | WWVTVYYGVPVWKEA | 237.1 |
|  | HLA-DRB1*07:01 | 33 | 47 |  | WWVTVYYGVPVWKEA | 359.8 |
|  | HLA-DPA1*01/DPB1*04:01 | 34 | 48 |  | WVTVYYGVPVWKEAT | 64.5 |
|  | HLA-DPA1*01:03/DPB1*02:01 | 34 | 48 |  | WVTVYYGVPVWKEAT | 50.4 |
|  | HLA-DPA1*02:01/DPB1*01:01 | 34 | 48 |  | WVTVYYGVPVWKEAT | 87.2 |
|  | HLA-DRB1*01:01 | 34 | 48 |  | WVTVYYGVPVWKEAT | 86.3 |
|  | HLA-DRB1*04:01 | 34 | 48 |  | WVTVYYGVPVWKEAT | 205.3 |
|  | HLA-DRB1*04:05 | 34 | 48 |  | WVTVYYGVPVWKEAT | 228.7 |
|  | HLA-DRB1*07:01 | 34 | 48 |  | WVTVYYGVPVWKEAT | 441 |
|  | HLA-DPA1*01/DPB1*04:01 | 35 | 49 |  | VTVYYGVPVWKEATT | 77.8 |
|  | HLA-DPA1*01:03/DPB1*02:01 | 35 | 49 |  | VTVYYGVPVWKEATT | 49.4 |
|  | HLA-DPA1*02:01/DPB1*01:01 | 35 | 49 |  | VTVYYGVPVWKEATT | 87.5 |
|  | HLA-DRB1*01:01 | 35 | 49 |  | VTVYYGVPVWKEATT | 61.4 |
|  | HLA-DRB1*04:01 | 35 | 49 |  | VTVYYGVPVWKEATT | 162.1 |
|  | HLA-DRB1*04:05 | 35 | 49 |  | VTVYYGVPVWKEATT | 231.9 |
|  | HLA-DPA1*01/DPB1*04:01 | 36 | 50 |  | TVYYGVPVWKEATTT | 105.3 |
|  | HLA-DPA1*01:03/DPB1*02:01 | 36 | 50 |  | TVYYGVPVWKEATTT | 80.5 |
|  | HLA-DPA1*02:01/DPB1*01:01 | 36 | 50 |  | TVYYGVPVWKEATTT | 134.7 |
|  | HLA-DRB1*01:01 | 36 | 50 |  | TVYYGVPVWKEATTT | 101 |
|  | HLA-DRB1*04:01 | 36 | 50 |  | TVYYGVPVWKEATTT | 261.3 |
|  | HLA-DRB1*04:05 | 36 | 50 |  | TVYYGVPVWKEATTT | 301.1 |
|  | HLA-DPA1*01/DPB1*04:01 | 37 | 51 |  | VYYGVPVWKEATTTL | 164.2 |
|  | HLA-DPA1*01:03/DPB1*02:01 | 37 | 51 |  | VYYGVPVWKEATTTL | 168.4 |
|  | HLA-DPA1*02:01/DPB1*01:01 | 37 | 51 |  | VYYGVPVWKEATTTL | 220.7 |
|  | HLA-DRB1*01:01 | 37 | 51 |  | VYYGVPVWKEATTTL | 165 |
|  | HLA-DRB1*04:05 | 37 | 51 |  | VYYGVPVWKEATTTL | 499.4 |
|  | HLA-DRB5*01:01 | 37 | 51 |  | VYYGVPVWKEATTTL | 261 |
|  | HLA-DPA1*01/DPB1*04:01 | 38 | 52 |  | YYGVPVWKEATTTLF | 326.1 |
|  | HLA-DPA1*01:03/DPB1*02:01 | 38 | 52 |  | YYGVPVWKEATTTLF | 288.8 |
|  | HLA-DRB1*01:01 | 38 | 52 |  | YYGVPVWKEATTTLF | 264.4 |
|  | HLA-DQA1*01:01/DQB1*05:01 | 17 | 31 | LLQYWSQEL | LKYLGNLLQYWSQEL | 251.9 |
|  | HLA-DRB1*15:01 | 17 | 31 |  | LKYLGNLLQYWSQEL | 12 |
|  | HLA-DQA1*01:01/DQB1*05:01 | 18 | 32 |  | KYLGNLLQYWSQELK | 230.8 |
|  | HLA-DRB1*01:01 | 18 | 32 |  | KYLGNLLQYWSQELK | 168.5 |
|  | HLA-DRB1*15:01 | 18 | 32 |  | KYLGNLLQYWSQELK | 9.5 |
|  | HLA-DRB4*01:01 | 18 | 32 |  | KYLGNLLQYWSQELK | 112.4 |
|  | HLA-DQA1*01:01/DQB1*05:01 | 19 | 33 |  | YLGNLLQYWSQELKN | 218.1 |
|  | HLA-DRB1*01:01 | 19 | 33 |  | YLGNLLQYWSQELKN | 120.4 |
|  | HLA-DRB1*04:05 | 19 | 33 |  | YLGNLLQYWSQELKN | 243.5 |
|  | HLA-DRB1*15:01 | 19 | 33 |  | YLGNLLQYWSQELKN | 8.6 |
|  | HLA-DRB4*01:01 | 19 | 33 |  | YLGNLLQYWSQELKN | 96.4 |
|  | HLA-DPA1*01:03/DPB1*02:01 | 20 | 34 |  | LGNLLQYWSQELKNS | 220.5 |
|  | HLA-DQA1*01:01/DQB1*05:01 | 20 | 34 |  | LGNLLQYWSQELKNS | 250.3 |
|  | HLA-DRB1*01:01 | 20 | 34 |  | LGNLLQYWSQELKNS | 73.4 |
|  | HLA-DRB1*04:05 | 20 | 34 |  | LGNLLQYWSQELKNS | 430.3 |
|  | HLA-DRB1*15:01 | 20 | 34 |  | LGNLLQYWSQELKNS | 12.5 |
|  | HLA-DRB4*01:01 | 20 | 34 |  | LGNLLQYWSQELKNS | 88.1 |
|  | HLA-DPA1*01:03/DPB1*02:01 | 21 | 35 |  | GNLLQYWSQELKNSA | 332.2 |
|  | HLA-DQA1*01:01/DQB1*05:01 | 21 | 35 |  | GNLLQYWSQELKNSA | 314.9 |
|  | HLA-DRB1*01:01 | 21 | 35 |  | GNLLQYWSQELKNSA | 184.4 |
|  | HLA-DRB1*15:01 | 21 | 35 |  | GNLLQYWSQELKNSA | 15 |
|  | HLA-DRB4*01:01 | 21 | 35 |  | GNLLQYWSQELKNSA | 114.3 |
|  | HLA-DPA1*01/DPB1*04:01 | 22 | 36 |  | NLLQYWSQELKNSAI | 444.9 |
|  | HLA-DPA1*01:03/DPB1*02:01 | 22 | 36 |  | NLLQYWSQELKNSAI | 298.8 |
|  | HLA-DQA1*01:01/DQB1*05:01 | 22 | 36 |  | NLLQYWSQELKNSAI | 400.3 |
|  | HLA-DRB1*01:01 | 22 | 36 |  | NLLQYWSQELKNSAI | 194.9 |
|  | HLA-DRB1*15:01 | 22 | 36 |  | NLLQYWSQELKNSAI | 20.9 |
|  | HLA-DPA1*01/DPB1*04:01 | 23 | 37 |  | LLQYWSQELKNSAIS | 412.6 |
|  | HLA-DPA1*01:03/DPB1*02:01 | 23 | 37 |  | LLQYWSQELKNSAIS | 448.4 |
|  | HLA-DRB1*15:01 | 23 | 37 |  | LLQYWSQELKNSAIS | 48.3 |
| P51 | HLADPA1*01/DPB1*04:01 | 11 | 25 | FLWMGYELH | HQKEPPFLWMGYELH | 68.1 |
|  | HLADPA1*01:03/DPB1*02:1 | 11 | 25 |  | HQKEPPFLWMGYELH | 35 |
|  | HLADPA1*03:01/DPB1*04:2 | 11 | 25 |  | HQKEPPFLWMGYELH | 95.3 |
|  | HLA-DRB1*01:01 | 11 | 25 |  | HQKEPPFLWMGYELH | 299.3 |
|  | HLA-DRB1*04:04 | 11 | 25 |  | HQKEPPFLWMGYELH | 356.3 |
|  | HLA-DRB1*04:05 | 11 | 25 |  | HQKEPPFLWMGYELH | 205.1 |
|  | HLADPA1*01/DPB1*04:01 | 12 | 26 |  | QKEPPFLWMGYELHP | 52.1 |
|  | HLADPA1*01:03/DPB1*02:1 | 12 | 26 |  | QKEPPFLWMGYELHP | 32 |
|  | HLADPA1*02:01/DPB1*01:1 | 12 | 26 |  | QKEPPFLWMGYELHP | 147.7 |
|  | HLADPA1*03:01/DPB1*04:2 | 12 | 26 |  | QKEPPFLWMGYELHP | 75.9 |
|  | HLA-DRB1*01:01 | 12 | 26 |  | QKEPPFLWMGYELHP | 315.1 |
|  | HLA-DRB1*04:04 | 12 | 26 |  | QKEPPFLWMGYELHP | 214.5 |
|  | HLA-DRB1*04:05 | 12 | 26 |  | QKEPPFLWMGYELHP | 203.9 |
|  | HLADPA1*01/DPB1*04:01 | 13 | 27 |  | KEPPFLWMGYELHPD | 44.1 |
|  | HLADPA1*01:03/DPB1*02:1 | 13 | 27 |  | KEPPFLWMGYELHPD | 32.4 |
|  | HLADPA1*02:01/DPB1*01:1 | 13 | 27 |  | KEPPFLWMGYELHPD | 124.4 |
|  | HLADPA1*03:01/DPB1*04:2 | 13 | 27 |  | KEPPFLWMGYELHPD | 62.2 |
|  | HLADQA1*01:01/DQB1*051 | 13 | 27 |  | KEPPFLWMGYELHPD | 474.9 |
|  | HLADQA1*05:01/DQ1 | 13 | 27 |  | KEPPFLWMGYELHPD | 458.9 |
|  | HLA-DRB1*01:01 | 13 | 27 |  | KEPPFLWMGYELHPD | 322.8 |
|  | HLA-DRB1*04:01 | 13 | 27 |  | KEPPFLWMGYELHPD | 423.9 |
|  | HLA-DRB1*04:04 | 13 | 27 |  | KEPPFLWMGYELHPD | 150.3 |
|  | HLA-DRB1*15:01 | 13 | 27 |  | KEPPFLWMGYELHPD | 277 |
|  | HLADPA1*01/DPB1*04:01 | 14 | 28 |  | EPPFLWMGYELHPDK | 44.1 |
|  | HLADPA1*01:03/DPB1*02:1 | 14 | 28 |  | EPPFLWMGYELHPDK | 36.3 |
|  | HLADPA1*02:01/DPB1*01:1 | 14 | 28 |  | EPPFLWMGYELHPDK | 118.3 |
|  | HLADPA1*03:01/DPB1*04:2 | 14 | 28 |  | EPPFLWMGYELHPDK | 52.1 |
|  | HLA-DRB1*01:01 | 14 | 28 |  | EPPFLWMGYELHPDK | 291.8 |
|  | HLA-DRB1*04:01 | 14 | 28 |  | EPPFLWMGYELHPDK | 325.2 |
|  | HLA-DRB1*04:04 | 14 | 28 |  | EPPFLWMGYELHPDK | 147.5 |
|  | HLA-DRB1*15:01 | 14 | 28 |  | EPPFLWMGYELHPDK | 242.6 |
|  | HLA-DRB5*01:01 | 14 | 28 |  | EPPFLWMGYELHPDK | 344.4 |
|  | HLADPA1*01/DPB1*04:01 | 15 | 29 |  | PPFLWMGYELHPDKW | 44.9 |
|  | HLADPA1*01:03/DPB1*02:1 | 15 | 29 |  | PPFLWMGYELHPDKW | 50.6 |
|  | HLADPA1*02:01/DPB1*01:1 | 15 | 29 |  | PPFLWMGYELHPDKW | 135.2 |
|  | HLADPA1*03:01/DPB1*04:2 | 15 | 29 |  | PPFLWMGYELHPDKW | 64 |
|  | HLA-DRB1*01:01 | 15 | 29 |  | PPFLWMGYELHPDKW | 346.6 |
|  | HLA-DRB1*04:01 | 15 | 29 |  | PPFLWMGYELHPDKW | 364.7 |
|  | HLA-DRB1*04:04 | 15 | 29 |  | PPFLWMGYELHPDKW | 134.4 |
|  | HLA-DRB1*15:01 | 15 | 29 |  | PPFLWMGYELHPDKW | 245.5 |
|  | HLA-DRB5*01:01 | 15 | 29 |  | PPFLWMGYELHPDKW | 381.6 |
|  | HLA- DPA1*01/DPB1*04:01 | 16 | 30 |  | PFLWMGYELHPDKWT | 68.3 |
|  | HLADPA1*01:03/DPB1*02:1 | 16 | 30 |  | PFLWMGYELHPDKWT | 100.1 |
|  | HLADQA1*03:01/DQB1*03:2 | 58 | 72 | WEFVNTPPL | ATWIPEWEFVNTPPL | 410.4 |
|  | HLA-DRB1*01:01 | 58 | 72 |  | ATWIPEWEFVNTPPL | 418.4 |
|  | HLA-DRB1*04:04 | 58 | 72 |  | ATWIPEWEFVNTPPL | 53.4 |
|  | HLA-DRB1*04:05 | 58 | 72 |  | ATWIPEWEFVNTPPL | 154.6 |
|  | HLA-DRB1*07:01 | 58 | 72 |  | ATWIPEWEFVNTPPL | 13.1 |
|  | HLA-DRB1*09:01 | 58 | 72 |  | ATWIPEWEFVNTPPL | 165.5 |
|  | HLA-DRB1*01:01 | 59 | 73 |  | TWIPEWEFVNTPPLV | 163 |
|  | HLA-DRB1*04:04 | 59 | 73 |  | TWIPEWEFVNTPPLV | 45.6 |
|  | HLA-DRB1*04:05 | 59 | 73 |  | TWIPEWEFVNTPPLV | 136.7 |
|  | HLA-DRB1*07:01 | 59 | 73 |  | TWIPEWEFVNTPPLV | 12.1 |
|  | HLA-DRB1*09:01 | 59 | 73 |  | TWIPEWEFVNTPPLV | 128.7 |
|  | HLA-DRB1*01:01 | 60 | 74 |  | WIPEWEFVNTPPLVK | 68.9 |
|  | HLA-DRB1*04:04 | 60 | 74 |  | WIPEWEFVNTPPLVK | 31.4 |
|  | HLA-DRB1*04:05 | 60 | 74 |  | WIPEWEFVNTPPLVK | 109.7 |
|  | HLA-DRB1*07:01 | 60 | 74 |  | WIPEWEFVNTPPLVK | 9.9 |
|  | HLA- DPA1*02:01/DPB1*01:01 | 60 | 74 |  | WIPEWEFVNTPPLVK | 456.3 |
|  | HLA- DPA1*02:01/DPB1*01:01 | 61 | 75 |  | IPEWEFVNTPPLVKL | 288.4 |
|  | HLA-DRB1*01:01 | 61 | 75 |  | IPEWEFVNTPPLVKL | 21.2 |
|  | HLA-DRB1*04:04 | 61 | 75 |  | IPEWEFVNTPPLVKL | 31.9 |
|  | HLA-DRB1*04:05 | 61 | 75 |  | IPEWEFVNTPPLVKL | 95.1 |
|  | HLA-DRB1*07:01 | 61 | 75 |  | IPEWEFVNTPPLVKL | 9.4 |
|  | HLA-DRB1*09:01 | 61 | 75 |  | IPEWEFVNTPPLVKL | 73.9 |
|  | HLA-DRB1*15:01 | 61 | 75 |  | IPEWEFVNTPPLVKL | 411.3 |
|  | HLA-DRB1*01:01 | 62 | 76 |  | PEWEFVNTPPLVKLW | 31.5 |
|  | HLA-DRB1*04:04 | 62 | 76 |  | PEWEFVNTPPLVKLW | 57.8 |
|  | HLA-DRB1*04:05 | 62 | 76 |  | PEWEFVNTPPLVKLW | 130.4 |
|  | HLA-DRB1*07:01 | 62 | 76 |  | PEWEFVNTPPLVKLW | 11.3 |
|  | HLA-DRB1*09:01 | 62 | 76 |  | PEWEFVNTPPLVKLW | 94 |
|  | HLA-DRB1*01:01 | 63 | 77 |  | EWEFVNTPPLVKLWY | 32.4 |
|  | HLA-DRB1*04:04 | 63 | 77 |  | EWEFVNTPPLVKLWY | 242.9 |
|  | HLA-DRB1*04:05 | 63 | 77 |  | EWEFVNTPPLVKLWY | 181.2 |
|  | HLA-DRB1*07:01 | 63 | 77 |  | EWEFVNTPPLVKLWY | 13.1 |
|  | HLA-DRB1*09:01 | 63 | 77 |  | EWEFVNTPPLVKLWY | 111.1 |
|  | HLA-DRB1*04:04 | 64 | 78 |  | WEFVNTPPLVKLWYQ | 349.4 |
|  | HLA-DRB1*04:05 | 64 | 78 |  | WEFVNTPPLVKLWYQ | 268.8 |
|  | HLA-DRB1*07:01 | 64 | 78 |  | WEFVNTPPLVKLWYQ | 15.3 |
|  | HLA-DRB1*01:01 | 64 | 78 |  | WEFVNTPPLVKLWYQ | 49.5 |
